# Quantification of multicellular colonization in tumor metastasis using exome sequencing data

**DOI:** 10.1101/542324

**Authors:** Jo Nishino, Shuichi Watanabe, Fuyuki Miya, Takashi Kamatani, Keith A Boroevich, Tatsuhiko Tsunoda

## Abstract

Metastasis is a major cause of cancer-related mortality, and it is essential to understand how metastasis occurs in order to overcome it. One relevant question is the origin of a metastatic tumor cell population. Although the hypothesis of a single-cell origin for metastasis from a primary tumor has long been prevalent, several recent studies using mouse models have supported a multi-cellular origin of metastasis. Human bulk whole-exome sequencing (WES) studies also have demonstrated a multiple ‘clonal’ origin of metastasis, with different mutational compositions. Specifically, there has not yet been strong research to determine how many founder cells colonize a metastatic tumor. To address this question, we developed a method to quantify the ‘founder cell population size’ in a metastasis using paired WES data from primary and metachronous metastatic tumors. Simulation studies demonstrated the proposed method gives unbiased results with sufficient accuracy in the range of realistic settings. Applying the proposed method to real WES data from four colorectal cancer patients, all samples supported a multi-cellular origin of metastasis and the founder size was quantified, ranging from 3 to 15 cells. Such a wide-ranging founder sizes estimated by the proposed method suggests that there are large variations in genetic similarity between primary and metastatic tumors in the same subjects, which might be involved in (dis)similarity of drug responses between tumors.

**Novelty and impact:** Applying our proposed method to the exome sequence data from four colorectal cancer patients, we showed the ‘multi-cellular origin’, not the classical ‘single-cell origin’, of metastasis is correct, with founder sizes quantified in the range of 3 to 15 cells. These wide-ranging founder sizes suggest large variation in genetic similarity between both tumors, which may affect the (dis)similarity of drug response in primary and metastatic tumors.

## Introduction

Metastasis is the main cause of cancer-related death. Although it is essential to understand its mechanisms and dynamics of distant site colonization in order to properly treat it, until recently little has been known. The founder cell population size of a metastatic tumor is one of the most important parameters for metastasis dynamics, which involves the change of mutational compositions from the primary to metastatic tumors (Figure 1). The drastic genetic changes in the metastatic tumor from the primary one, brought by the limited cell migration, i.e., ‘bottleneck effect’, might result in a difference in drug response between both tumors in the same patients.

**Figure 1.**
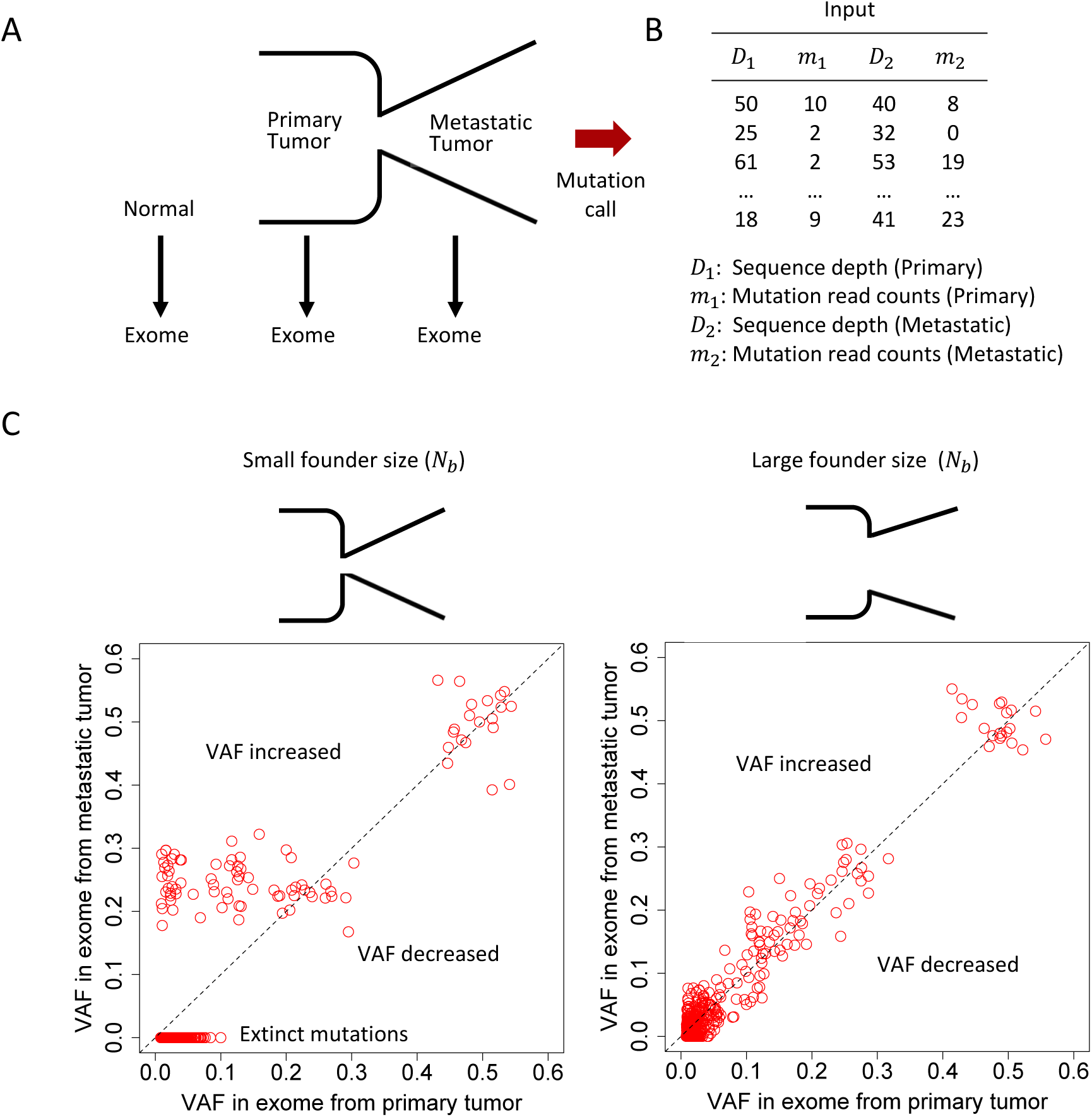
A schematic view of the proposed methodology. (A) Exome data from paired primary and metastatic tumors, and normal tissue. (B) Input of the method. (C) Illustration of basic premise for the estimation of founder sizes by computer simulations. Low correlation of observed VAFs in exome between the primary and the metastatic tumors in the small founder size, *N*_*b*_ =2 (left). High correlation of observed VAFs between the primary and metastatic tumors in the large founder size, *N*_*b*_=50 (right).

Although the hypothesis that a metastatic tumor originates from a single tumor cell has been long prevalent ^1-3^, several recent studies using mouse models of cancer demonstrated multicellular seeding ^4-6^. In humans, bulk whole-exome sequencing (WES) studies of metastatic tumors, often including primary tumors from the same individuals, demonstrated metastases to have originated from multiple clones, where a ‘clone’ was a cluster of tumor cells belonging to the same phylogenetic clade estimated by the variant allele frequency information ^7, 8^. While founder ‘cells’, but not ‘clones’, in the metastatic tumor have another clear meaning in understanding metastatic dynamics, the quantification of multicellular colonization has not been attempted so far in human metastatic tumors.

Here, we propose a method to quantify the founder cell population size of a metastatic tumor using a paired WES data from the primary and metachronous metastatic tumors. The method uses the outputs from commonly used mutation callers, i.e., variant allele frequencies (mutant allele counts and sequence depths), and quickly estimates the founder size unbiasedly in a realistic range. We applied our proposed method to the high-depth WES data from a study for four colorectal cancer (CRC) patients.

## Methods

### Overview for quantifying founder cell population size in metastasis

We use paired WES data of a primary and metachronous metastatic tumors together with the data from the normal tissue (Figure 1A). The input file is composed of sequence depths, *D*_1_ and *D*_2_, and the mutation read counts, *m*_1_ and *m*_2_ for each called mutation in the primary and metastatic tumors, respectively (Table 1 and Figure 1B; See supplementary Appendix and supplementary Figure S1 for more details of the input file). When the founder population size is large, the variant allele frequencies (VAFs) for called mutations in the metastatic show high similarity to those in the primary tumor (Figure 1C). Conversely, when the size of founder cells is small, the VAFs in the primary and metastatic tumors are not so correlated (Figure 1C). In this case, due to the severe ‘bottleneck effect’, many variants can become extinct or have a significantly higher VAF in the metastatic tumor.

**Table 1.** Notations in the Model and the simulation study.

| Notation | Description |
| --- | --- |
| $N_b$ | Founder cell population size, to be estimated. |
| $R$ | Number of mutations with or more than $m_{1(min)}$ mutation reads in the primary tumor. |
| $m_1, m_2$ | Mutation read counts for the primary ($m_1$) and metastatic tumors ($m_2$). |
| $m_{1(min)}$ | Minimum mutation read count in WES data from the primary tumor. For estimating $N_b$, we use only the sites with or more than $m_{1(min)}$ mutant reads. |
| $D_1, D_2$ | Sequence depths for the primary ($D_1$) and metastatic tumors ($D_2$). |
| $p_1, p_2$ | Population VAFs in the primary ($p_1$) and metastatic tumor ($p_2$). |
| $f(p_1)$ | Probability density of $p_1$. |
| $M_b$ | Number of mutant cells among $N_b$ founders. |
| $\gamma_1, \gamma_2$ | Tumor purity in the WES samples from the primary ($\gamma_1$) and metastatic tumors ($\gamma_2$). |
| Additional notations in the simulation study. |  |
| $K$ | Number of clonal mutations in the initial primary tumor. |
| $\mu$ | Mutation rate per tumor-cell division in the primary tumor. |
| $N_1$ | Cell population size in the final primary tumor. |
| $\bar{D}$ | Mean sequence depth in the primary and metastatic tumor. |
| $\gamma$ | Tumor purity in the WES samples from the primary and metastatic tumors ($\gamma_1=\gamma_2$). |

### Model and estimation methods

Consider a diploid tumor cell population in a primary tumor. One somatic variant in the population has the VAF, *p*_1_, or the cancer cell fraction (CCF), 2*p*_1_ (see Table 1 for notations). The models assume no recurrence mutation at the same sites and therefore the VAF is at most 0.5, *p*_1_ ≤ 0.5. The VAF follows some distribution, *p*_1_ ∼ *f*(*p*_1_), as is properly assumed in the present implementation assuming the ‘neutral’ evolution with high cell birth rate for tumor population^9, 10^ (see Implementation section in supplementary Appendix; and see Results section for the robustness of the assumptions). In bulk-WES of the primary tumor, the sampled mutation read count, *m*_1_, at the variant site with sequence depth, *D*_1_, follows a binomial distribution with parameters, *D*_1_ and *p*_1_,

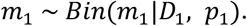

In bulk-WES of the metastatic tumor, the sampled mutation read count, *m*_2_, at the variant site with sequence depth, *D*_2_, is generated by a composite process of metastatic colonization and exome sequencing as follows:

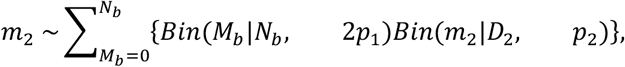

where the *N*_*b*_, *M*_*b*_ and *p*_2_ are the number of founder cells (founder population size) in metastatic colonization, the number of mutant cells in the *N*_*b*_ founder cells, and the VAF in the metastatic tumor, respectively. In the above distribution for *m*_2_, the *N*_*b*_ founder cells are randomly selected from the primary tumor and colonize a metastatic site. The *M*_*b*_ mutant cells in the metastatic site follows a binomial distribution with parameters *N*_*b*_ and 2*p*_1_ (mutant cell fraction). The sampled mutation read count, *m*_2_, follows a binomial distribution with parameter *D*_2_ and *p*_2_, where *p*_2_ is given by 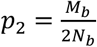.

Taken together, the probability of observing *m*_1_ and *m*_2_ mutations in the primary and metastatic exome with depths *D*_1_ and *D*_2_, respectively, is given by

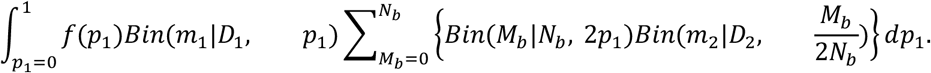

For quality control, we use only the sites with or more than *m*_1(*min*)_ (>0) mutant reads in the primary tumor. Note that, in the metastatic tumor, all mutations called in the primary tumor are tracked in order to use greater information on VAF change from the primary to the metastatic tumor. Finally, the probability of observing *m*_1_(≥ *m*_1(*min*)_) and *m*_2_(≥ 0) mutation reads in the primary and metastatic tumors, respectively, is expressed as

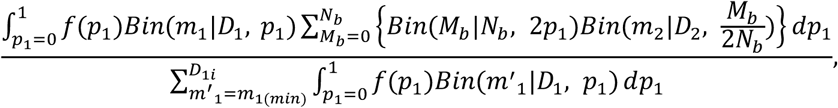

where *m*′_1_ is possible read counts in the primary tumor. Explicitly, let *p*_1*i*_, *D*_1*i*_, *m*_1*i*_, *D*_2*i*_ and *m*_2*i*_ denote *p*_1_, *D*_1_, *m*_1_, *D*_2_ and *m*_2_ for the specific *i*-th variant site, respectively. Assuming independencies among all *R* variants, each with *m*_1*i*_(≥ *m*_1(*min*)_) mutation reads in the metastatic tumor, the likelihood of the founder size, *N*_*b*_, is given by

*Likelihood*(*N*_*b*_)

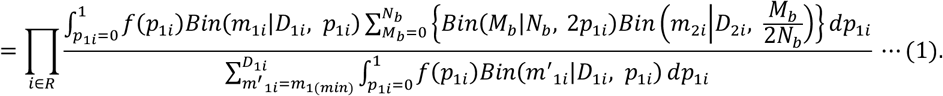

By maximizing the likelihood (1), we obtain the maximum likelihood estimate of *N*_*b*_ (for implementation details, see supplementary Appendix). In reality, the independence assumption among variants does not hold since the unit of the tumor evolution is the cell, and mutations in the same cell evolve and colonize a metastatic site together. The effect of the independence assumption on the estimation of *N*_*b*_ is investigated below using simulations.

The tumor purities, the fraction of cancer cells, in the primary (*γ*_1_) and metastatic tumor tissue samples (*γ*_2_), are incorporated into the model simply by replacing *p*_1*i*_ in the term *Bin*(*m*_1*i*_|*D*_1*i*_, *p*_1*i*_) with *γ*_1_*p*_1*i*_ and 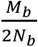 in the term 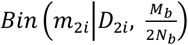 with 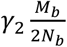, respectively.

## Results

### Validation of our proposed method by simulations

#### Pure birth model

We assessed our proposed method using simulated data, generated by a ‘pure birth model’ for tumor evolution (see Methods and supplementary Appendix for details; see also Table 1 for notations). Briefly, a single tumor cell with *K* mutations generates two daughter cells, each with average *μ* new mutations, and cell divisions repeat until the population has grown to the final primary tumor size, *N*_1_. *N*_*b*_ cells from the *N*_1_ cells make up a metastatic tumor. Exome samples in the primary and metastatic tumor have depth 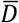 and purity *γ*. Our proposed method was applied to the sites with ≥ *m*_1(*min*)_ mutant reads in the primary tumor. We ran 100 simulation for each parameter sets. Mouse models have suggested that metastasis occurs via colonization of one CTC cluster rather than serial arrivals of CTC clusters (or single CTCs) and that the most CTC clusters contain between 2 to 20 tumor cells, with median of 6 ^6^. We mainly focused on this range of founder sizes in the simulations.

In Figure 2A-D, all simulations were performed under the conditions of *N*_1_ = 100,000, *μ*=2.5. Firstly, the effect of varying mean depth, 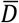, on the estimation of *N*_*b*_, was investigated under *K* =50, *γ* =1, and *m*_1(*min*)_ =2 (Figure 2A). The number of variants generated in the exome samples in the simulations were realistic, ranging from one to around five hundred (Supplementary Figure S2B). In cases of *N*_*b*_=2, 5, 10 and 20, when 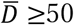, the medians of estimates were very close to the true values, i.e., the estimator is median-unbiased, and the estimation accuracy is good. For example, when the depth was 50, the medians of the estimates (and interquartile ranges; IQRs) were 5.0 (4.0, 6.0), 10.0 (8.0, 13.0), and 20.0 (15.0, 28.25) for the true *N*_*b*_=5, 10 and 20, respectively. The unbiasedness with 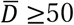 held for larger *N*_*b*_ (for *N*_*b*_ =1-100, see supplementary Figure S2B). The estimation accuracy got better as sequence depth increased. Even when the depth was 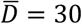, the precision and accuracy were acceptable, the medians of estimates (IQRs) were 5.0 (5.0, 6.0), 12.0 (8.0, 18.25), and 21.0 (14.0, 35.0) for the true *N*_*b*_ =5, 10 and 20, respectively. Under 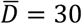, particularly for larger *N*_*b*_ ≥ 30, *N*_*b*_ was biasedly estimated and a reliable estimation was difficult to obtain (for *N*_*b*_=1-100, see supplementary Figure S2A). Note that, for all depth settings, the relative estimation errors were better for smaller *N*_*b*_, as you can see from the smaller log-scaled boxplots of the estimated *N*_*b*_ in Figure 1A (see also supplementary Figure S2A).

**Figure 2.**
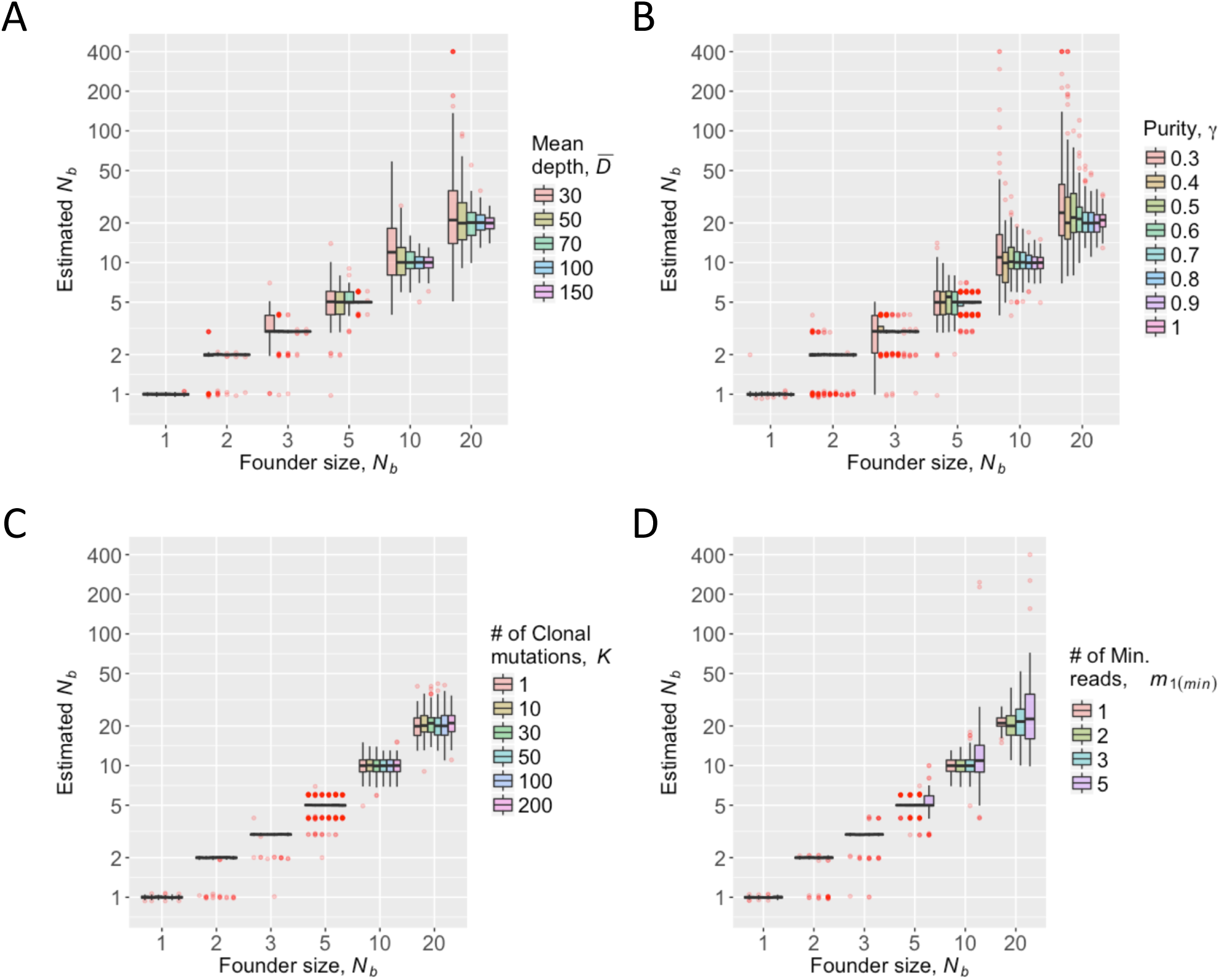
Valid quantification of founder size, *N*_*b*_, confirmed by simulations. All simulations used the same primary tumor population size, *N*_1_ *=* 100,000, and mutation rate per cell division per exome, *µ* = 2.5. (A) Varying mean sequencing depth, 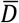 for *K*=50, *γ* =1, and *m*_1(*min*)_=2. (B) Varying tumor purity, *γ*, for 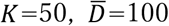, and *m*_1(*min*)_=2. (C) Varying number of clonal mutations, *K*, for 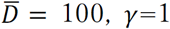 and *m*_1(*min*)_=2. (D) Varying minimum number of mutation reads, *m*_1(*min*)_, for 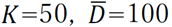, and *γ* =1. (Variants with *m*_1(*min*)_ or more mutation reads were used.)

Next, the effects of the tumor purity, *γ*, on the estimation were investigated under 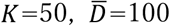, and *m*_1(*min*)_=2 (Figure 2B). When *γ* ≥ 50%, the estimation was median-unbiased and the accuracy was acceptable. The medians of the estimates (IQRs) were 5.0 (5.0, 6.0), 10.0 (8.0, 12.0), and 20.0 (15.0, 28.0) for the true *N*_*b*_=5, 10 and 20, respectively. In conjunction with the result of Figure 1A, defining the ‘effective sequence depth’ as the depth multiplied by tumor purity, the proposed method gave unbiased results with acceptable accuracy when the effective sequence depth was 50. In the case of less purity, and large founder size e.g., *γ* ≤ 40% and *N*_*b*_ ≥ 30, a reliable estimation was difficult obtain (for *N*_*b*_=1-100, see supplementary Figure S3).

In the algorithm for *N*_*b*_ estimation, the proportion of clonal mutations in the primary tumors are fixed at 10%. However, true clonal mutations vary among tumors. Thus, the impacts of the number of clonal mutations were investigated under 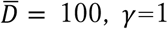 and *m*_1(*min*)_=2 (Figure 2C). The number of clonal mutations in the population, *K*, had no effect on both the unbiasedness and the accuracy of estimation of *N*_*b*_. The same is true for larger *N*_*b*_ with various number of variants in WES samples (for *N*_*b*_ =1-100, see supplementary Figure S4).

As input of the proposed method, we use variants with *m*_1(*min*)_ or more mutation reads in the primary tumor. Then, the effects of various values of *m*_1(*min*)_ on the estimation of *N*_*b*_ were investigated under 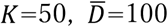, and *γ*=1 (Figure 2D). The estimation results for up to *N*_*b*_=100 were also assessed (supplementary Figure S5). In the case of *m*_1(*min*)_ = 5, the estimation accuracy was worse than those for *m*_1(*min*)_ < 5. The worse accuracy was not due to lower numbers of variants used for input (for the case of larger number of variants, see supplementary Figure S6 replacing *µ*=2.5 with *µ*=12.5). For the case of including singletons in the input (*m*_1(*min*)_ = 1), a small upward bias can occur (for more clear bias in the large *N*_*b*_, see supplementary Figure S5). Thus, we recommend the criteria of ‘at least 2 or 3 mutation read counts’, *m*_1(*min*)_=2 or 3, for the input of the proposed method.

Simulations were performed mainly under the conditions of the primary tumor size, *N*_1_ = 100,000 and mutation rate, *µ*=2.5. When values of *N*_1_ ranging from 1,000 to 300,000 were used under 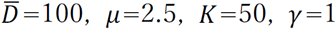, and *m*_1(*min*)_ =2, the behavior of estimates were generally the same as that under *N*_1_ = 100,000 (supplementary Figure S7). When values of *µ* ranging from 0.5 to 10 were used under 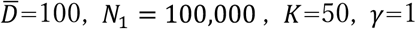, and *m*_1(*min*)_ =2, the behavior of estimates were generally the same as that under *µ* = 2.5 although the estimation accuracy was a little lower as the mutation rate is small (supplementary Figure S8).

#### Robustness for cell death and selection

So far, in the development of primary tumor, it was assumed there was no cell death and no difference in cell division rates. The violation of the assumptions might make estimation of *N*_*b*_ difficult, since VAF distribution, *f*(*p*_1_), possibly become different from the postulated distribution under ‘neutral’ evolution with high cell birth rate of tumor population. Here, we investigated the consequences of the violation, keeping the other settings as Figure 1, i.e., *µ* =2.5, *K* =50, *γ*=1, and 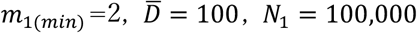. We ran 100 simulation for each parameter set.

Firstly, to investigate the effect of ‘cell death’, death rate, *d*, and, birth rate, *b*, per unit time were introduced. Limiting the case to *d* < *b*, which means steady growth of tumor population, various values, *d*=0.01, 0.1 and 0.2, against unit birth rate, *b* = 1, were assumed (the ratio of *d* to *b* define the evolutional system). For any death rates, *d*=0.01, 0.1 and 0.2, the estimator for the founder size, *N*_*b*_, is median-unbiased and the estimation accuracy is sufficient, as with the case of no death (*d*=0) (supplementary Figure S9A). This is due to the fact that VAF distribution for *d* ≠ 0 is not so far that for no death case as long as *d* ≪ *b* ^10^ (supplementary Figure S9B-D).

Secondly, we considered the case that one positively selective subclone in the primary tumor appears in the WES samples^11^. One starting primary tumor cell with *b* = 1, *d* = 0.1 are assumed to evolve and at the time when the population reach to a particular population size, *N*_*sub*.*occ*._, one selectively advantageous mutation occurs. The subclone with the advantageous mutation have large birth rate, *b*=2, 5 or 10. The values of *N*_*sub*.*occ*._ are determined so that corresponding frequencies of the selective mutation are low (∼2%), middle (∼16%) and high (∼30%) at the WES sampling point. Although distributions of VAF were shifted to the frequency of the selective mutation (supplementary Figure S10-12B, C, D), the estimator for the founder size, *N*_*b*_, is median-unbiased and the estimation accuracy is sufficient, as well as for case of no selective subclone (supplementary Figure S10-12A).

Lastly, we considered the case that many mutations with small effects are accumulated in the developmental process of the primary tumor. Neutral mutations and non-neutral mutations occur with the probabilities of 0.3 and 0.7, respectively, which mimics synonymous and non-synonymous mutation rates in the exon region. The birth rate of a cell acquiring a new non-neutral mutation (*b*_*new*_) is given by multiplying the present birth rate (*b*_*pre*_) by (1+ *a*), i.e., *b*_*new*_ = *b*_*pre*_(1+ *a*), where *a* is a coefficient for birth rate and set as *a* = ±0.01, ±0.05, ±0.1, ±0.15 and ± 0.2. Positive and negative value of *a* mean deleterious and advantageous mutations, respectively. The death rate is always set to be one-tenth of population mean of birth rates. Advantageous mutations, particularly when *a* ≥ 0.1, shift VAF distribution toward intermediate frequency (supplementary Figure S13B-K). When deleterious or advantageous mutations with *a* ≤0.1, the estimator for the founder size, *N*_*b*_, is median-unbiased and the estimation accuracy is sufficient, as with the case of no selection, *a* = 0 (supplementary Figure S13A). When strong selection with *a* ≥ 0.15, the estimator is biased upward and the accuracy is not so good. But it is unrealistic that one-third of all mutations would have such strong effects.

#### Real data analysis

We used the high-depth WES data from a study for four colorectal cancer (CRC) patients, which included at least one primary and metachronous metastatic tumor sample per patient ^8^. For each patient, the metastatic tumor(s) were sampled 1-3 years after the removal of the primary tumor(s). Called mutation summary information for each tumor were derived from the article ^8^. Our method was applied to the four paired primary and metastatic tumors’ data in the four patients, the data from Pri-1 and Met-1 for each patient (the primary and metastatic tumors were arbitrarily labeled Pri-x and Met-x, respectively, per patient). We conducted quality-controls and used the called mutations satisfying the following criteria: within 1,000 sequencing depths in the primary and metastatic tumors, having at least two mutation reads in the primary tumor, i.e., *m*_1(*min*)_=2, having no mutation read in the normal sample in the primary and metastatic tumors, and without no copy number aberrations. The last criterion ensured diploid tumor populations, which is assumed in the current model, and copy number aberrations were retrieved from the article ^8^.

The proposed method was applied to the mutations that passed the quality-controls using the estimated tumor purities by PurBayes ^12^. Mutations considered to be possible errors, many of which had higher VAFs (supplementary Figure S14) were removed. We then performed the definitive analyses to estimate the founder population size of metastatic tumors. The average number of variants used ranged from 79-195 for the four samples, with average sequence depths of 83.4-146.7 and 83.4-146.7 in the primary and metastatic tumor exomes, respectively (Table 2). The tumor purities were 0.23−0.72 and 0.29-0.85 in the primary and metastatic tumor samples, respectively (Table 2). The observed VAFs looked to be somewhat correlated in the primary and metastatic tumor in each patient (supplementary Figure S15). Estimated founder population sizes (80% confidence intervals) were estimated to be 15 (13.0, 17.0), 3 (2.0, 4.0), 8 (6.0, 9.0), and 14 (9.0, 21.1) for subject A01, A02, A03, A04, respectively (Figure 3). Consistent results were obtained when the same analyses were carried out using all variations without limiting diploid regions (Table 2 for the mutation summary, supplementary Figure S14 for removed outliers, supplementary Figure S15 for VAFs, and Figure 3 for the estimated founder sizes).

**Table 2.** Summary of WES data for CRC subjects from Wei et al. (2017).

| Subject | # of SNV (R) | Tumor | Mean Depth | Purity (95%CI) |
| --- | --- | --- | --- | --- |
| A01<br>- Diploid | 195 | Pri-1 (colon) | 137.6 | 0.46 (0.44, 0.48) |
|  |  | Met-1 (liver) | 153.1 | 0.48 (0.47, 0.50) |
| - All | 233 | Pri-1 (colon) | 138.4 | 0.46 (0.44, 0.48) |
|  |  | Met-1 (liver) | 154.4 | 0.48 (0.46, 0.49) |
| A02<br>- Diploid | 165 | Pri-1 (colon) | 130.3 | 0.23 (0.22, 0.24) |
|  |  | Met-1 (lung) | 129.7 | 0.29 (0.28, 0.30) |
| - All | 209 | Pri-1 (colon) | 139.5 | 0.45 (0.43, 0.48) |
|  |  | Met-1 (lung) | 141.1 | 0.29 (0.28, 0.30) |
| A03<br>- Diploid | 72 | Pri-1 (colon) | 83.4 | 0.25 (0.23, 0.26) |
|  |  | Met-1 (liver) | 102.6 | 0.59 (0.58, 0.61) |
| - All | 110 | Pri-1 (colon) | 87.4 | 0.46 (0.43, 0.49) |
|  |  | Met-1 (liver) | 105.2 | 0.80 (0.76, 0.85) |
| A04<br>- Diploid | 193 | Pri-1 (colon) | 146.7 | 0.72 (0.68, 0.76) |
|  |  | Met-1 (lung) | 148.2 | 0.85 (0.81, 0.89) |
| - All | 224 | Pri-1 (colon) | 149.6 | 0.70 (0.65, 0.74) |
|  |  | Met-1 (lung) | 150.0 | 0.84 (0.80, 0.88) |

Purities and 95% credible intervals (CIs) were estimated by PurBayes [9]. Diploid: Results using only diploid regions (excluding copy number aberrations). All: Results using all exome regions (including copy number aberrations).

**Figure 3.**
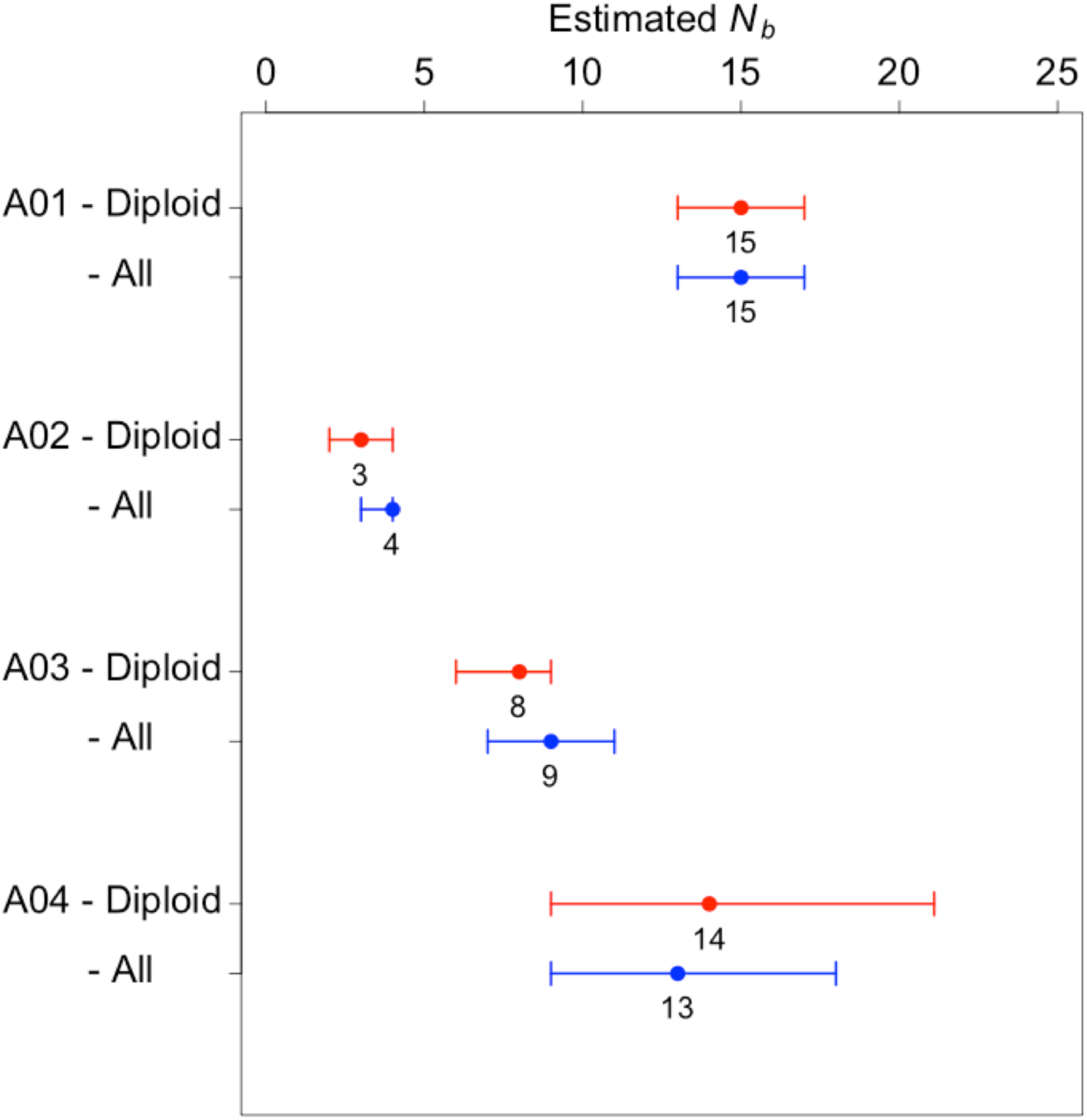
Estimated founder sizes (*N*_*b*_) for the four colorectal cancer reported by Wei et al. (2017). Results using only diploid regions (excluding copy number aberrations) are shown in red. Results using all exome regions (including copy number aberrations) are shown in blue. Circles with bars indicate maximum likelihood estimates of *N*_*b*_ and these 80% confidence intervals, based on 100 non-parametric bootstrap samples.

## Discussion

We developed a method to quantify the founder population size in metastasis using a paired WES data from primary and metachronous metastatic tumors. This method, implicitly using the fact that higher (lower) genetic similarity between the primary and metastatic tumors results from a larger (smaller) founder size (Figure 1C), enables us to unbiasedly estimate the founder population size with sufficient accuracy in the range of realistic founder size and settings, e.g., sequencing depth, purity and number of variations (Figure 2 and supplementary Figure S1-7). The method is also robust to the realistic model of primary tumor evolution, including cell death and several selective differences in cell birth (supplementary Figure S9-13). Although relative estimation errors become worse as the founder size became larger, this weakness is overcome by deeper sequencing, i.e., WES data with × 150 depth give sufficient accuracy even for the founder size 100 (supplementary Figure S1). The proposed method also shows the advantage of using VAF information (mutation read counts and depths) rather than using only the presence or absence of mutations, to infer the tumor evolutionary process, as has been applied so far ^9, 11, 13^.

In real data analysis of four colorectal cancer patients, our method supported the multi-cellular origin of metastatic tumors, which is consistent with the observation of recent mouse model studies ^4-6^ and the suggestion from WES studies ^7, 8^. Our method further quantified the founder population sizes ranging 3 to 15 cells for CRC subjects ^8^. The wide-ranging founder size in metastasis might result in large variations of genetic similarity between primary and metastatic tumors, which might cause variation in drug responses between primary and metastatic tumors. In particular, when the founder population size is small, variants with drastically increased VAFs in the metastatic tumors might lead to difficulty in treatment.

In the context of population genetics, demographic history is a confounding factor for detecting and quantifying natural selection acting on the genome ^14, 15^. The same should be true for the evolution of a tumor population. A potential advantage of the proposed method is to identify selectively recruited mutations in the metastatic tumors under the inferred demographic model for tumor populations, i.e., the estimated founder size.

The limitations of our method are that it does not consider the time between the first exome sampling and metastatic occurrence and the time between metastatic occurrence and the second exome sampling. Particularly, in latter time periods, genetic drift in a small population of new metastatic tumor might not negligible. Our model does not distinguish between such genetic drift and the bottleneck effect of metastatic colonization, and the estimate of ‘the founder size’ reflects both these effects. A method that distinguishes both effects will be future work. In addition, there are possibly more complex cell migration patterns than our model, including reseeding or multisource seeding ^16, 17^, which are beyond the present study but worth investigating.

## Data Availability Statement

The data that support the findings of this study were derived from Supplementary Table5-12 in the reference number 8.

## Disclosure

The authors have declared no conflicts of interest.

The earlier version of this manuscript has been posted on a preprint server, bioRxiv (doi: doi.org/10.1101/542324)

## Abbreviations

WES: whole-exome sequencing
CRC: colorectal cancer
VAF: variant allele frequency
IQR: interquartile range

## References

1. Fidler IJ, Talmadge JE. Evidence that intravenously derived murine pulmonary melanoma metastases can originate from the expansion of a single tumor cell. Cancer research 1986;46: 5167–71.

2. Maddipati R, Stanger BZ. Pancreatic Cancer Metastases Harbor Evidence of Polyclonality. Cancer Discov 2015;5: 1086–97.

3. Talmadge JE, Fidler I. Evidence for the clonal origin of spontaneous metastases. Science 1982;217: 361–3.

4. Aceto N, Bardia A, Miyamoto DT, Donaldson MC, Wittner BS, Spencer JA, Yu M, Pely A, Engstrom A, Zhu H. Circulating tumor cell clusters are oligoclonal precursors of breast cancer metastasis. Cell 2014;158: 1110–22.

5. Cheung KJ, Ewald AJ. A collective route to metastasis: Seeding by tumor cell clusters. Science 2016;352: 167–9.

6. Cheung KJ, Padmanaban V, Silvestri V, Schipper K, Cohen JD, Fairchild AN, Gorin MA, Verdone JE, Pienta KJ, Bader JS, Ewald AJ. Polyclonal breast cancer metastases arise from collective dissemination of keratin 14-expressing tumor cell clusters. Proc Natl Acad Sci U S A 2016;113: E854–63.

7. Gundem G, Van Loo P, Kremeyer B, Alexandrov LB, Tubio JMC, Papaemmanuil E, Brewer DS, Kallio HML, Hognas G, Annala M, Kivinummi K, Goody V, et al. The evolutionary history of lethal metastatic prostate cancer. Nature 2015;520: 353–7.

8. Wei Q, Ye Z, Zhong X, Li L, Wang C, Myers RE, Palazzo JP, Fortuna D, Yan A, Waldman SA, Chen X, Posey JA, et al. Multiregion whole-exome sequencing of matched primary and metastatic tumors revealed genomic heterogeneity and suggested polyclonal seeding in colorectal cancer metastasis. Ann Oncol 2017;28: 2135–41.

9. Williams MJ, Werner B, Barnes CP, Graham TA, Sottoriva A. Identification of neutral tumor evolution across cancer types. Nat Genet 2016;48: 238–44.

10. Ohtsuki H, Innan H. Forward and backward evolutionary processes and allele frequency spectrum in a cancer cell population. Theor Popul Biol 2017;117: 43–50.

11. Williams MJ, Werner B, Heide T, Curtis C, Barnes CP, Sottoriva A, Graham TA. Quantification of subclonal selection in cancer from bulk sequencing data. Nat Genet 2018;50: 895–903.

12. Larson NB, Fridley BL. PurBayes: estimating tumor cellularity and subclonality in next-generation sequencing data. Bioinformatics 2013;29: 1888–9.

13. Sun R, Hu Z, Sottoriva A, Graham TA, Harpak A, Ma Z, Fischer JM, Shibata D, Curtis C. Between-region genetic divergence reflects the mode and tempo of tumor evolution. Nat Genet 2017;49: 1015–24.

14. Bamshad M, Wooding SP. Signatures of natural selection in the human genome. Nat Rev Genet 2003;4: 99–111.

15. Nielsen R, Hellmann I, Hubisz M, Bustamante C, Clark AG. Recent and ongoing selection in the human genome. Nat Rev Genet 2007;8: 857–68.

16. El-Kebir M, Satas G, Raphael BJ. Inferring parsimonious migration histories for metastatic cancers. Nat Genet 2018;50: 718–26.

17. Sanborn JZ, Chung J, Purdom E, Wang NJ, Kakavand H, Wilmott JS, Butler T, Thompson JF, Mann GJ, Haydu LE, Saw RP, Busam KJ, et al. Phylogenetic analyses of melanoma reveal complex patterns of metastatic dissemination. Proc Natl Acad Sci U S A 2015;112: 10995–1000.

