## supplementary Note for "Quantification of multicellular colonization in tumor metastasis using exome sequencing data"

### supplementary Appendix

**Preparation of input**

The input of the proposed methods is paired information for called somatic mutations in the primary and metachronous metastatic tumors, sequencing depths and mutation read counts. The following shows how to prepare the input data with GATK Mutect2 [1]. An example flow for preparing input data is shown in supplementary Figure 1.

Somatic mutations are called using Mutetct2 for the primary and metastatic tumors. In the call process, "--initial-tumor-lod 0" and "--tumor-lod-to-emit 0" should be set as Mutetct2 augments so that all possible mutations with small tumor LOD (log-odds) will be output. Then, as quality control for mutations, we use only mutations with $m_{1(min)}$ or more mutation reads in the primary tumor as input for our method. Note that all variant sites that passed the QC in the primary tumor are examined in the metastatic tumor, even in the case of few or no mutation reads, as was modeled in the Model section. When additional quality filtering processes are done for called mutations in the primary or metastatic tumors, we should take care to avoid sampling bias for VAF (or mutation read counts) since we take into account only the bias caused by the mutation detection criteria, $m_{1(min)}$. Note that using FilterMutectCalls, a dedicated filter of Mutect2, with default settings may discard many mutations with close to $m_{1(min)}$ mutation reads because of the low number of mutation reads. In order to resolve this, the --tumor-lod option should be set so that the "LOD threshold for calling tumor variant" is a small value.

**Implementation**

In order to maximize the likelihood given by (1) in the main body, we approximate the distribution of VAFs ($p_{1i}s$) in the primary tumor population, $f(p_{1i})$, as a discrete distribution having finite $n$ points of $p_{1i}$ with probabilities $\pi_1, \pi_2, \dots, \pi_n$. Then, the integrals with respective to $p_{1i}$ in (1) can be calculated by the corresponding summations. The $\pi_n$ specially represents the proportion of "clonal mutations", i.e., the proportion of mutations with CCF=1 or $p_{1i}$=0.5 (all tumor cells in heterozygous status). The other $\pi_j$s ($j = 1, \dots, n-1$) are the proportions of "subclonal" mutations with particular VAFs $(0 < p_{1i} < 0.5)$ in the population.

In preliminary simulation, taking into account of tumor evolution and sequencing, we found that small $p_{1i}$ (low-VAF) regions should be densely discretized for accurate estimation probably because the most mutations have low frequency in tumor, e.g., see [2, 3]. At the same time, using a large $n$ for dense discretization of $p_{1i}$ increases the computational burden. Therefore, we employed a discrete distribution having $n = 100$ values of $p_{1i}$, spaced evenly on a log-scale, to prioritize small $p_{1i}$ region. The first point of $p_1$ is set sufficiently small, $p_{1i} = 10^{-6}$. The cumulative discrete probabilities from 0.5 to the given values, $p_1$, are set to be ones from the theoretical expectation under the neutral evolution, i.e., $\propto (1/p_{1i} - 2)$ [3]. The proportion of clonal mutations, $\pi_n$, were assumed to be 10%. This arbitrarily fixed proportion worked well in the various true $\pi_n$ values (see the Results section).

We obtained the maximum likelihood estimate (MLE) of $N_b$ that maximized the likelihood given in (1) as follows. For $N_b$>10, we found the value of $N_b$ giving the maximum likelihood in (1) using a derivative-free optimization method, the golden section search (GSS) method [4]. For small $N_b$<10, we fully searched all values giving the maximum likelihood in (1) because of the possible non-concave function. Then, we determined MLE of $N_b$ combining the results from the GGS and full search methods. We applied a non-parametric bootstrap method to obtain the confidence intervals of MLE of $N_b$.

### Simulation model for validation of our proposed method

#### Pure birth model

For validation of the proposed method, we conducted computer simulations that mimic the data generation by tumor evolution and WES of the primary and metachronous metastatic tumors. The tumor evolution model was a "pure birth process", which is a special case of commonly used "birth-death process" for tumor evolution model [2, 3, 5]. Let us assume one diploid tumor cell with a particular number of clonal mutations, $K$, in a primary tumor generated two daughter cells with some new mutations, the number of which followed Poisson distributions with a particular mean, $\mu$. Next, a randomly chosen cell of the two generated two more daughter cells with some new mutations in the same manner. This process was repeated until the tumor grown up to the population of $N_1$ cells. At that time, each $N_1$ cell had a set of mutations and the $i$-th mutation had a frequency, $p_{1i}$, in the population. In the WES data from the primary tumor, the mutation read counts at the $i$-th mutation site, $m_{1i}$, followed a binomial distribution with parameters of sequence depth, $D_{1i}$, and VAF multiplied by the tumor purity, $(p_{1i} \times \gamma_1)$, where $D_{1i}$ followed a Poisson distribution with a parameter, $\bar{D}_1$. The information, $D_{1i}$ and $m_{1i}$, for the variants with $m_{1(min)}$ or more mutation read counts were recorded. Then, the $N_b$ founder cells were randomly selected from the $N_1$ primary tumor cells. A metastatic tumor that grew up without any selection or genetic drift would have the same VAFs as those of the founder cells. All mutations recorded in the primary tumor were examined in the sequence data from the metastatic tumor. Similarly to the case of the primary tumor, for the $i$-th mutation site, the mutation read counts $m_{2i}$, followed a binomial distribution with each sequence depth, $D_{2i}$, and VAF multiplied by the tumor purity, $(p_{2i} \times \gamma_2)$, where $D_2$ followed a Poisson distribution with a parameter, $\bar{D}_2$. Finally, using the dataset synthesized as above, $N_b$ was estimated by our proposed method. For simplicity, $\bar{D} = \bar{D}_1 = \bar{D}_2$ and $\gamma = \gamma_1 = \gamma_2$ were assumed.

**Model with death and selection**

In real tumor evolution, cell death and competition (selection) among tumor cells might occur [2, 3, 5]. As is described in the main body, by introducing birth rate and death rate, we investigated the effects of cell-death, one positively selective subclone, and many accumulated mutations with small effects in the primary tumor, on the estimate of $N_b$. At an arbitrary population size in the primary tumor, a subsequent event (birth or death) in the tumor population occurs in proportion to their birth and death rates, i.e., birth (or death) for a particular cell occurs with the probability of [birth (or death) rate for the cell]/[sum of all birth and death rates], until the tumor grown up to the population of $N_1$ cells. Other process regarding WES samples in the primary and metastatic tumors, and mutation in the daughter cells are the same as the pure birth model described above.

### References

1. Cibulskis K, Lawrence MS, Carter SL et al. Sensitive detection of somatic point mutations in impure and heterogeneous cancer samples. Nat Biotechnol 2013; 31: 213-219.
2. Ohtsuki H, Innan H. Forward and backward evolutionary processes and allele frequency spectrum in a cancer cell population. Theor Popul Biol 2017; 117: 43-50.
3. Williams MJ, Werner B, Barnes CP et al. Identification of neutral tumor evolution across cancer types. Nat Genet 2016; 48: 238-244.
4. Kiefer J. Sequential minimax search for a maximum. Proceedings of the American mathematical society 1953; 4: 502-506.
5. Williams MJ, Werner B, Heide T et al. Quantification of subclonal selection in cancer from bulk sequencing data. Nat Genet 2018; 50: 895-903.
