## supplementary Figures for "Quantification of multicellular colonization in tumor metastasis using exome sequencing data"

supplementary Figure S1.

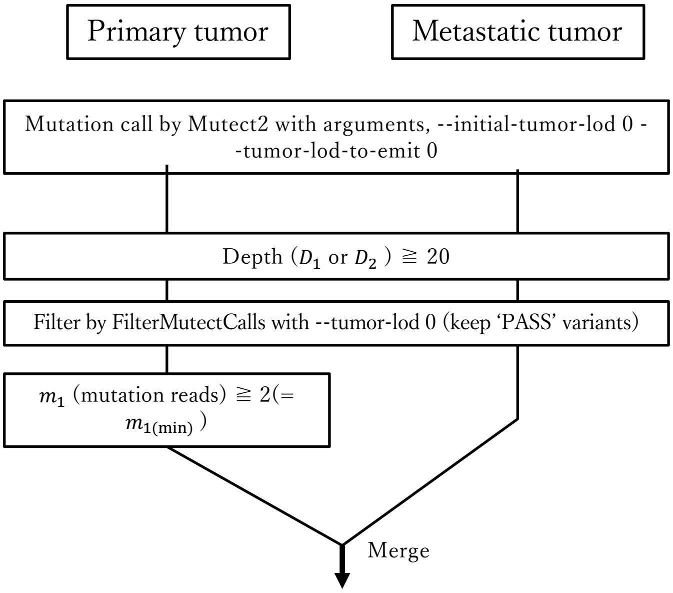


Input

| $D_1$ | $m_1$ | $D_2$ | $m_2$ |
|---|---|---|---|
| 50 | 10 | 40 | 8 |
| 25 | 2 | 32 | 0 |
| 61 | 2 | 53 | 19 |
| 33 | 2 | 28 | 0 |
| … | … | … | … |
| 18 | 9 | 41 | 23 |

supplementary Figure S1. Detailed example of the preparation of input data.

supplementary Figure S2.

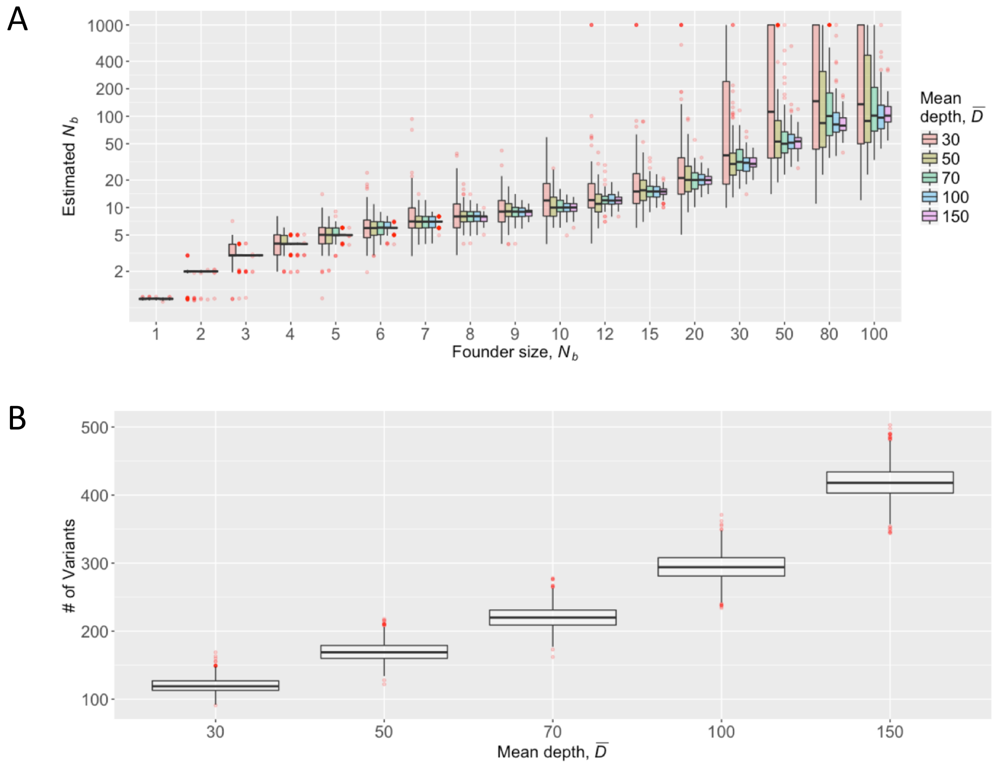


supplementary Figure S2. Quantification of founder size, $N_b$, using simulated data. The figure shows results of varying mean depth, $D$, and corresponds to Fig. 2A in the main body. Parameter values are the same as those of Fig. 2A, covering wider values of founder size, $N_b$. (A) Estimates of $N_b$. (B) Number of variants used for the estimation.

supplementary Figure S3.

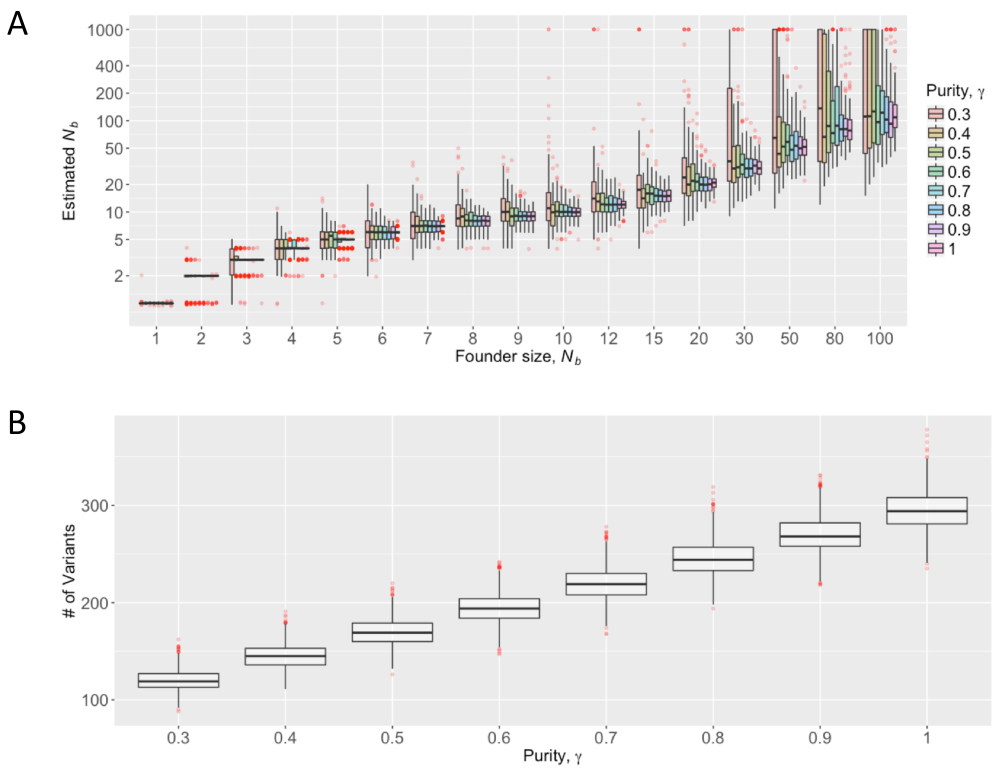


supplementary Figure S3. Quantification of founder size using simulated data. The figure shows results of varying purity, γ, and corresponds to Fig. 2B in the main body. Used parameter values are the same as those of Fig. 2B, covering wider values of founder size, $N_b$. (A) Estimates of $N_b$. (B) Number of variants used for the estimation.

supplementary Figure S4.

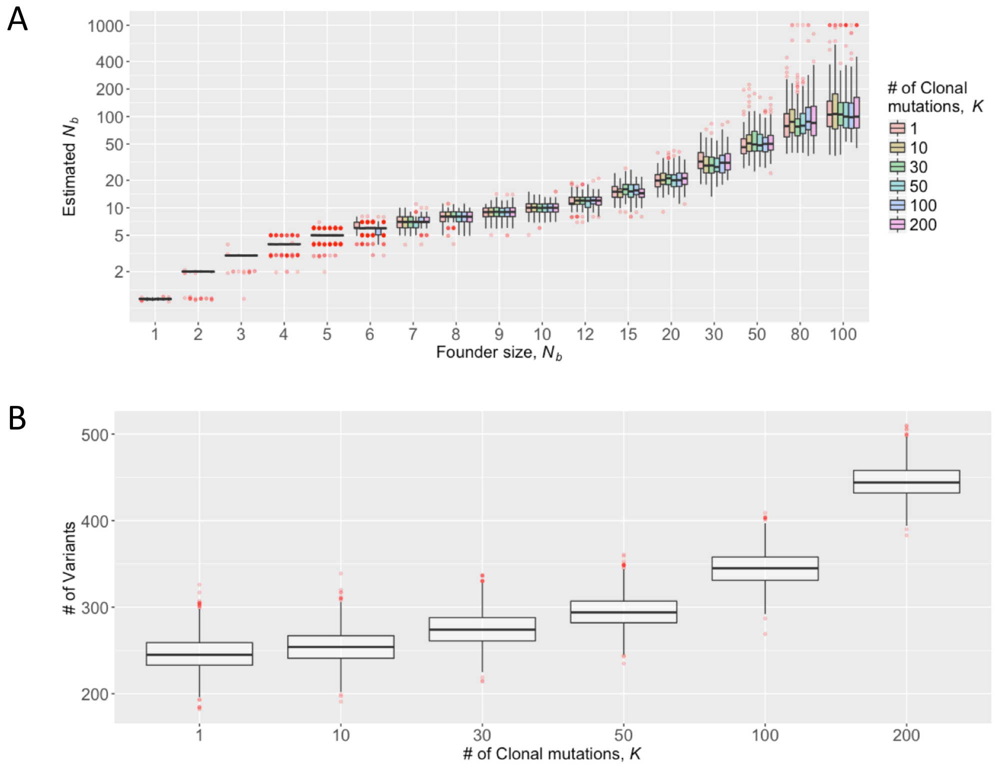


supplementary Figure S4. Quantification of founder size using simulated data. The figure shows results of varying number of clonal mutations, $K$, and corresponds to Fig. 2C in the main body. Used parameter values are the same as those of Fig. 2C, covering wider values of founder size, $N_b$. (A) Estimates of $N_b$. (B) Number of variants used for the estimation.

supplementary Figure S5.

A
B

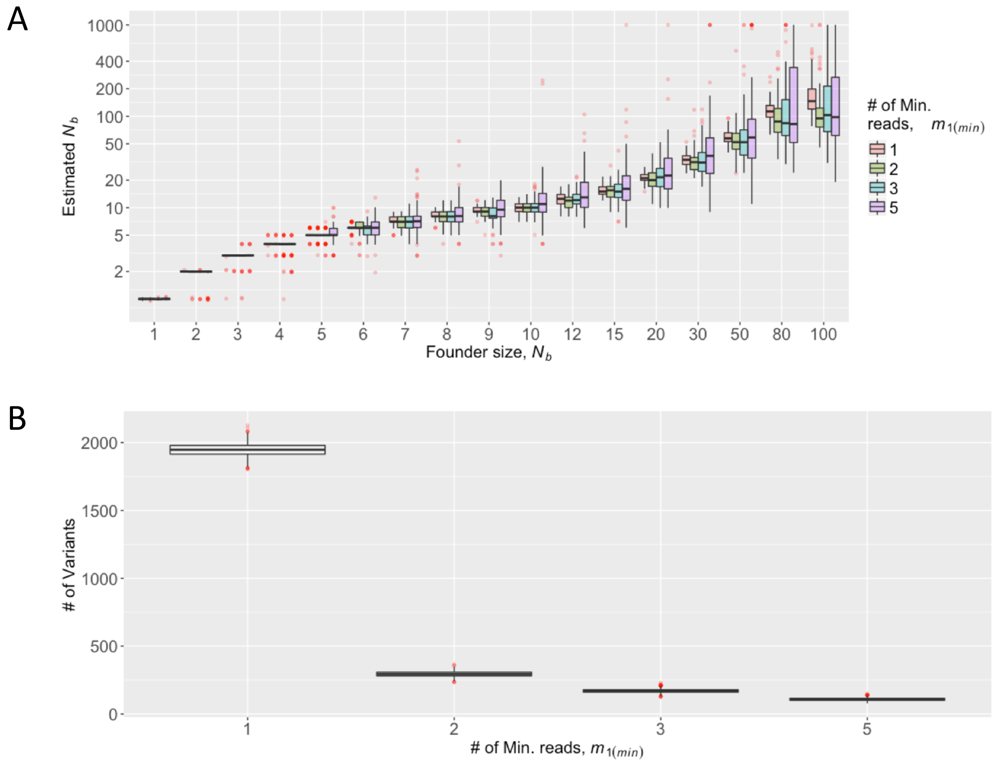


supplementary Figure S5. Quantification of founder size using simulated data. The figure shows results of varying minimum number of mutation reads, $m_{1(min)}$, and corresponds to Fig. 2D in the main body. Used parameter values are the same as those of Fig. 2D, covering wider values of founder size, $N_b$. (A) Estimates of $N_b$. (B) Number of variants used for the estimation.

supplementary Figure S6.

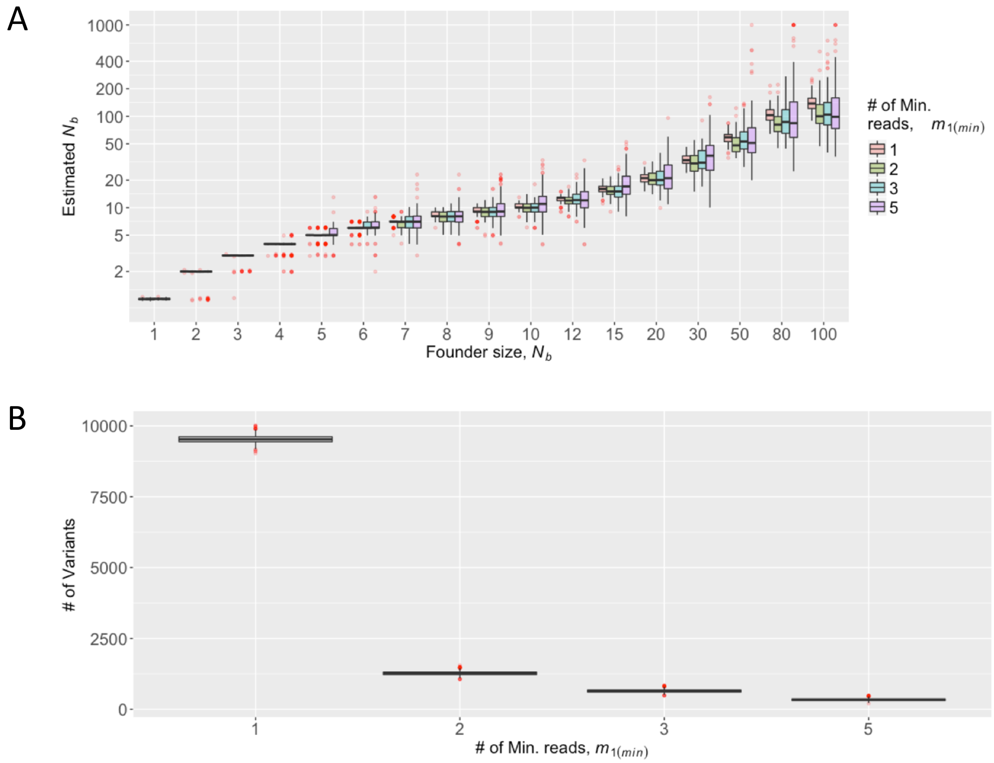


supplementary Figure S6. Quantification of founder size using simulated data. The figure shows results of varying minimum number of mutation reads, $m_{1(min)}$, and another version of Supplementary Figure S5 replacing $\mu$=5 with $\mu$=25. Other parameter values are the same as those of Fig. 2D, except for founder size, $N_b$. (A) Estimates of $N_b$. (B) Number of variants used for the estimation.

supplementary Figure S7.

A

B

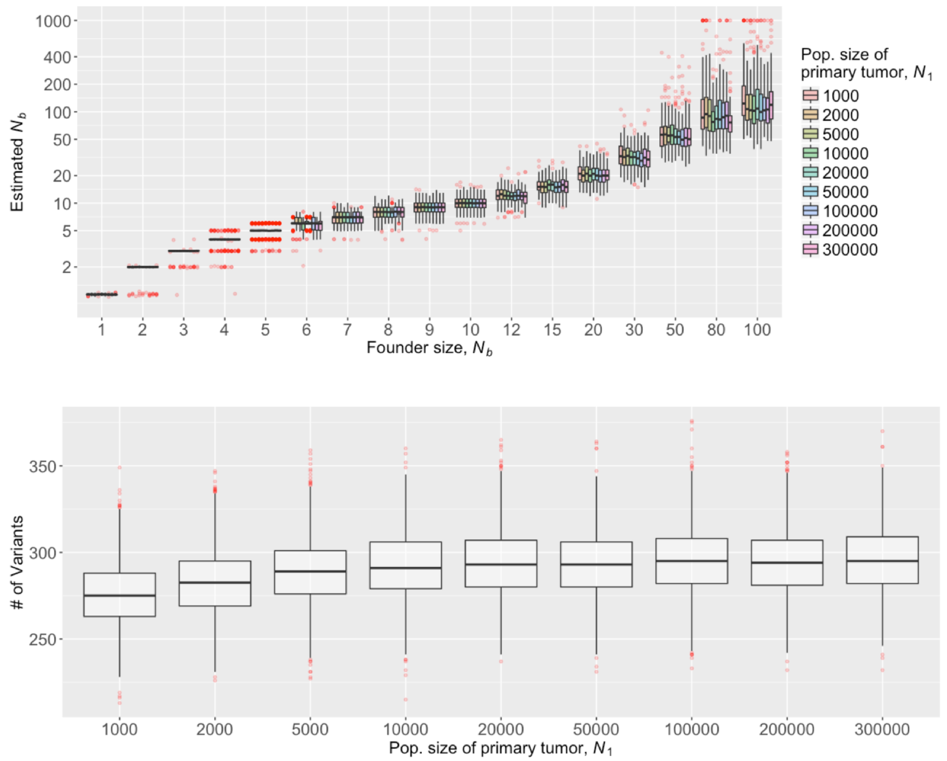


supplementary Figure S7. Quantification of founder size using simulated data. Results under varying primary tumor population size, $N_1$, $D$=100, $\mu$=2.5, $K$=50, $\gamma$=1, and $m_{1(min)}$=2. (A) Estimates of $N_b$. (B) Number of variants used for the estimation.

supplementary Figure S8.

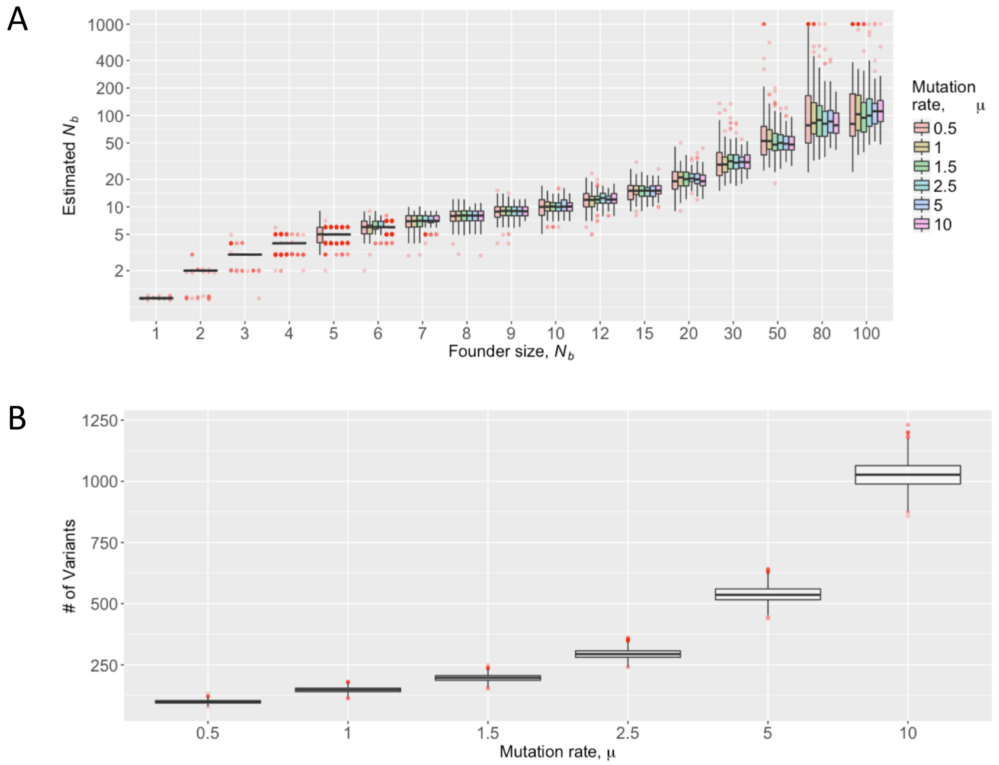


supplementary Figure S8. Quantification of founder size using simulated data. Results under varying mutation rates, $\mu$, $D$=100, $K$=50, $\gamma$=1, and $m_{1(min)}$=2. (A) Estimates of $N_b$. (B) Number of variants used for the estimation.

supplementary Figure S9.

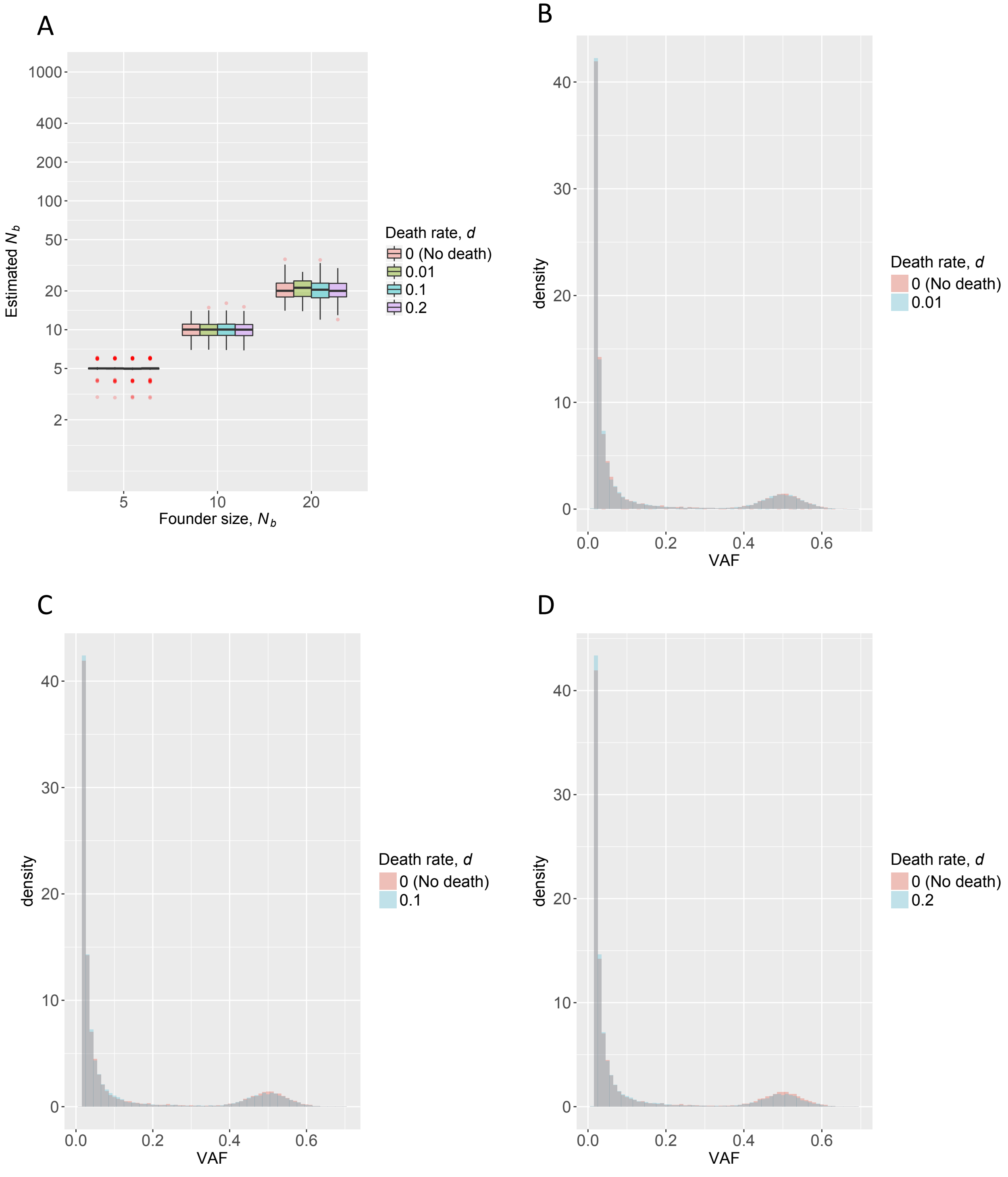


supplementary Figure S9. Quantification of founder size, $N_b$, using simulated data. The figure shows effect of cell death in the primary tumor evolution. Apart from death rate, $d$, parameter values are the same as Figure 1 in the main body, i.e., $\mu$=2.5, $K$=50, $\gamma$=1, and $m_{1(min)}$=2, $D$ = 100, $N_1$ = 100,000. (A) Estimates of $N_b$. (B)-(D) VAF distribution in the primary tumor WES.

supplementary Figure S10. ($b$=2)

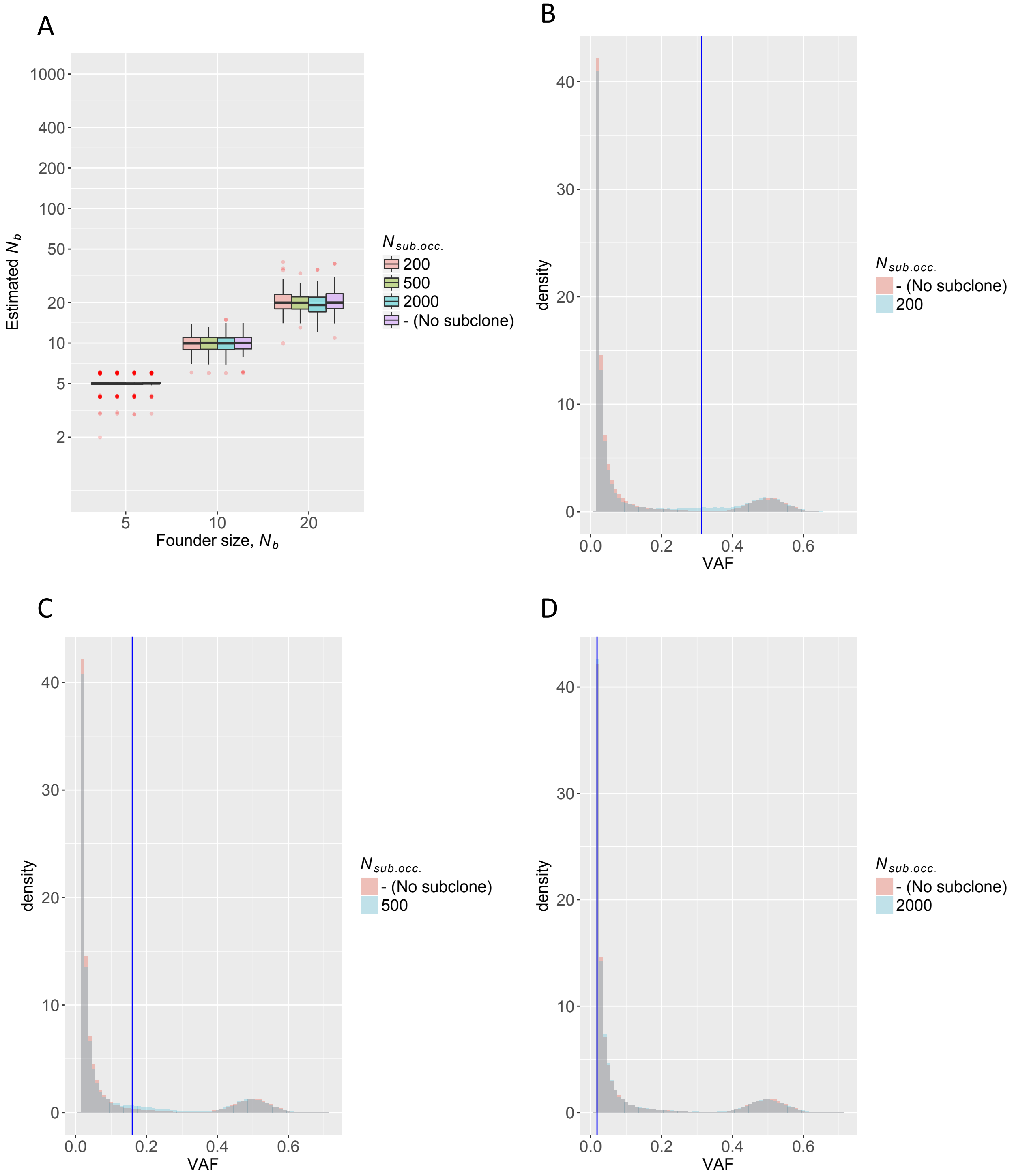


supplementary Figure S10. Quantification of founder size, $N_b$, using simulated data. The figure shows effect of one selective subclone with the birth rate, $b$=2, in the primary tumor. Apart from $N_{sub.occ.}$ (the population size of primary tumor at which one selective mutation occur), parameter values are the same as Figure 1 in the main body, i.e., $\mu$=2.5, $K$=50, $\gamma$=1, and $m_{1(min)}$=2, $D = 100$, $N_1 = 100{,}000$. (A) Estimates of $N_b$. (B)-(D) VAF distribution in the primary tumor WES. Blue vertical lines show theoretical expectation of VAF for the selective mutation.

supplementary Figure S11. ($b$=5)

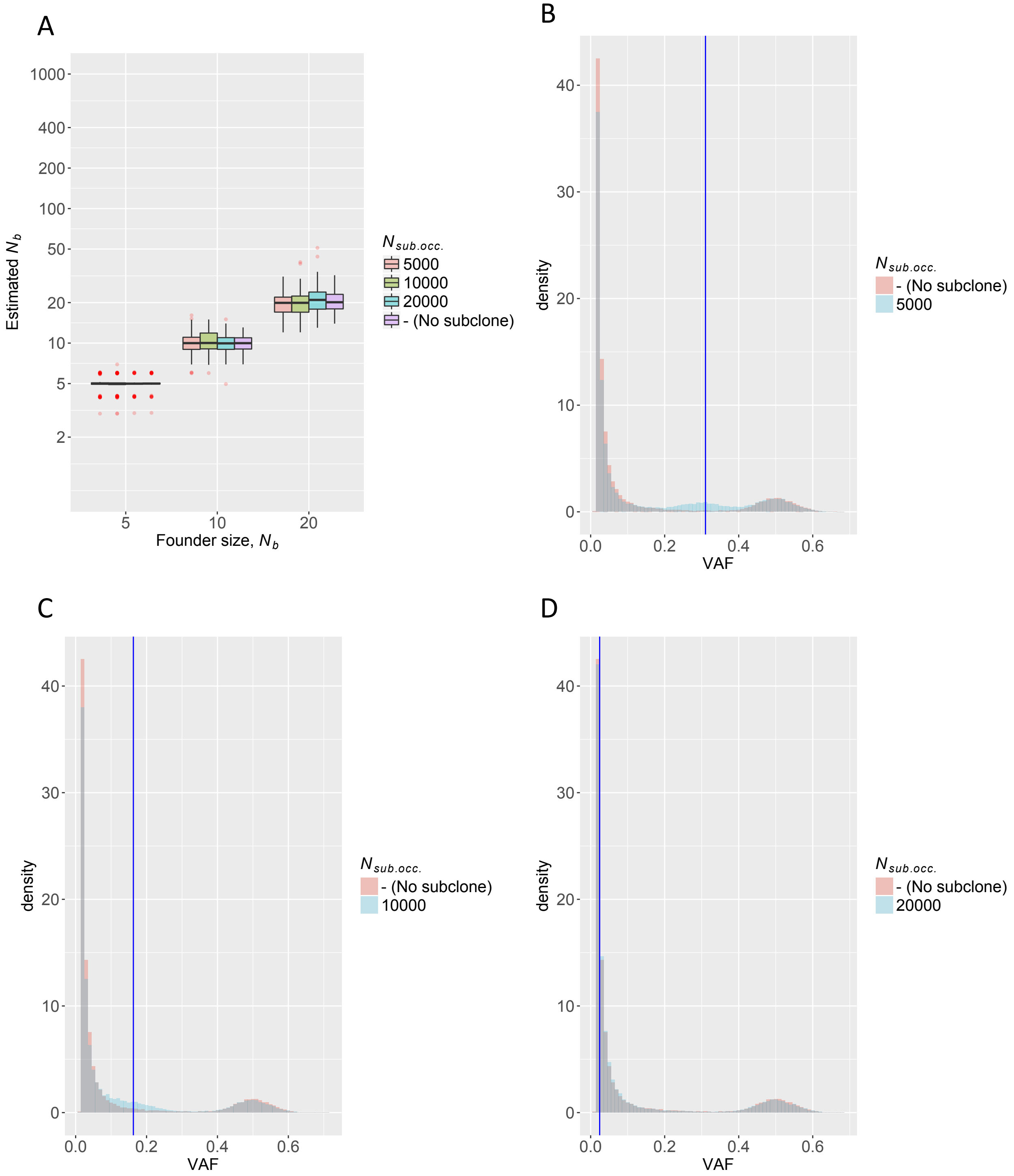


supplementary Figure S11. Quantification of founder size, $N_b$, using simulated data. The figure shows effect of one selective subclone with the birth rate, $b$=5, in the primary tumor. Apart from $N_{sub.occ.}$ (the population size of primary tumor at which one selective mutation occur), parameter values are the same as Figure 1 in the main body, i.e., $\mu$=2.5, $K$=50, $\gamma$=1, and $m_{1(min)}$=2, $D = 100$, $N_1 = 100{,}000$. (A) Estimates of $N_b$. (B)-(D) VAF distribution in the primary tumor WES. Blue vertical lines show theoretical expectation of VAF for the selective mutation.

supplementary Figure S12. ($b$=10)

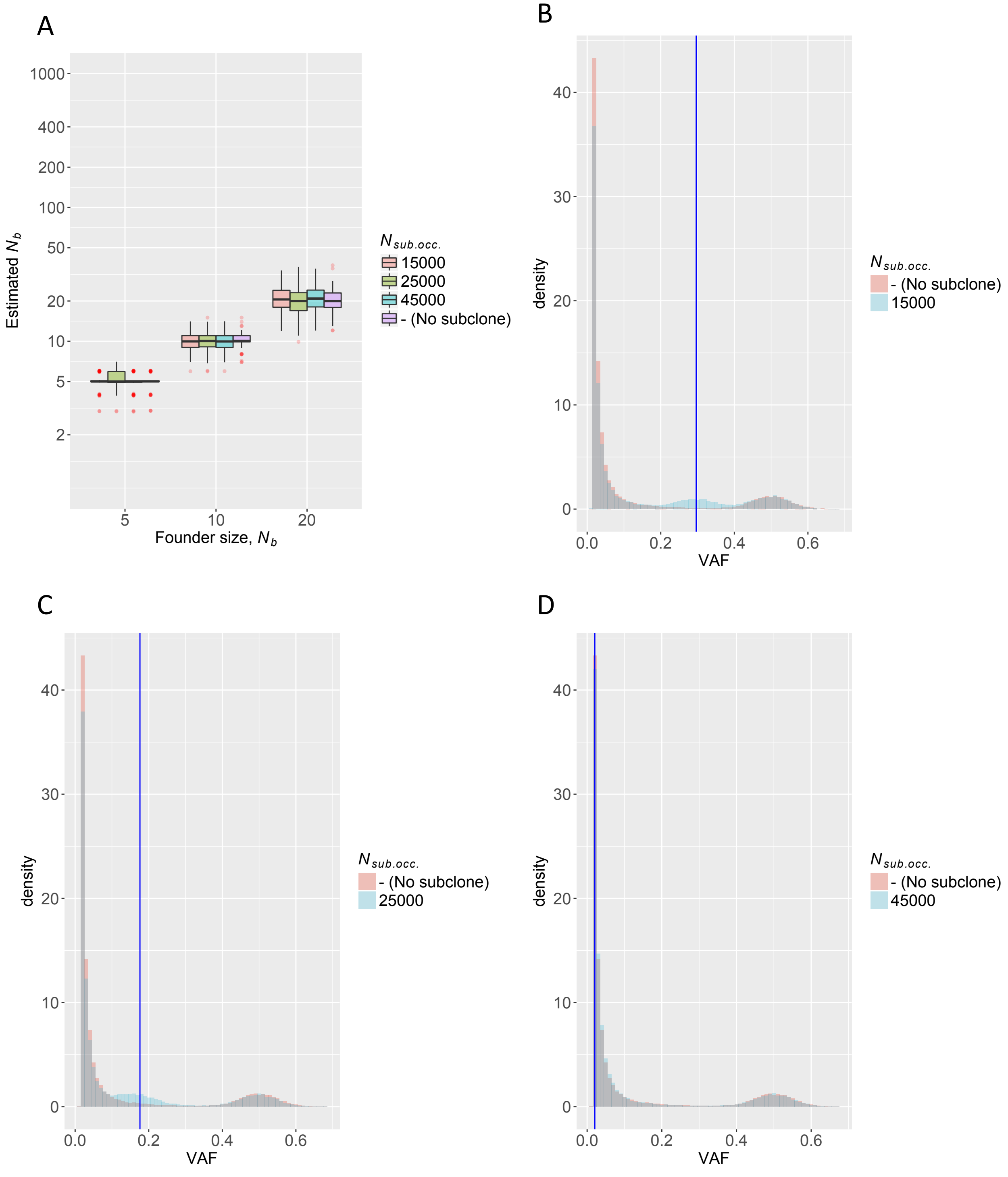


supplementary Figure S12. Quantification of founder size, $N_b$, using simulated data. The figure shows effect of one selective subclone with the birth rate, $b$=10, in the primary tumor. Apart from $N_{sub.occ.}$ (the population size of primary tumor at which one selective mutation occur), parameter values are the same as Figure 1 in the main body, i.e., $\mu$=2.5, $K$=50, $\gamma$=1, and $m_{1(min)}$=2, $D$ = 100, $N_1$ = 100,000. (A) Estimates of $N_b$. (B)-(D) VAF distribution in the primary tumor WES. Blue vertical lines show theoretical expectation of VAF for the selective mutation.

supplementary Figure S13.

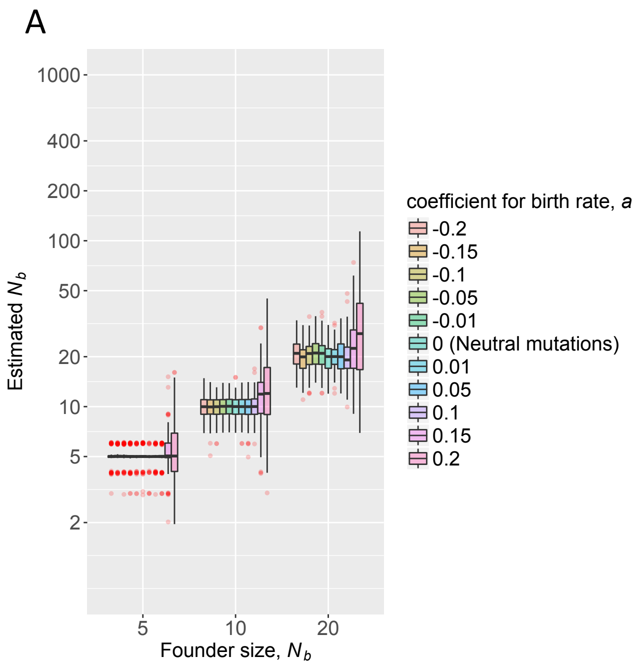


supplementary Figure S13. Quantification of founder size, $N_b$, using simulated data. The figure shows effect of many accumulated mutations with small effects in the primary tumor. Apart from a coefficient for birth rate, $a$, parameter values are the same as Figure 1 in the main body, i.e., $\mu$=2.5, $K$=50, $\gamma$=1, and $m_{1(min)}$=2, $D$ = 100, $N_1$ = 100,000. (A) Estimates of $N_b$. (B)-(K) VAF distribution in the primary tumor WES.

supplementary Figure S13. Continued

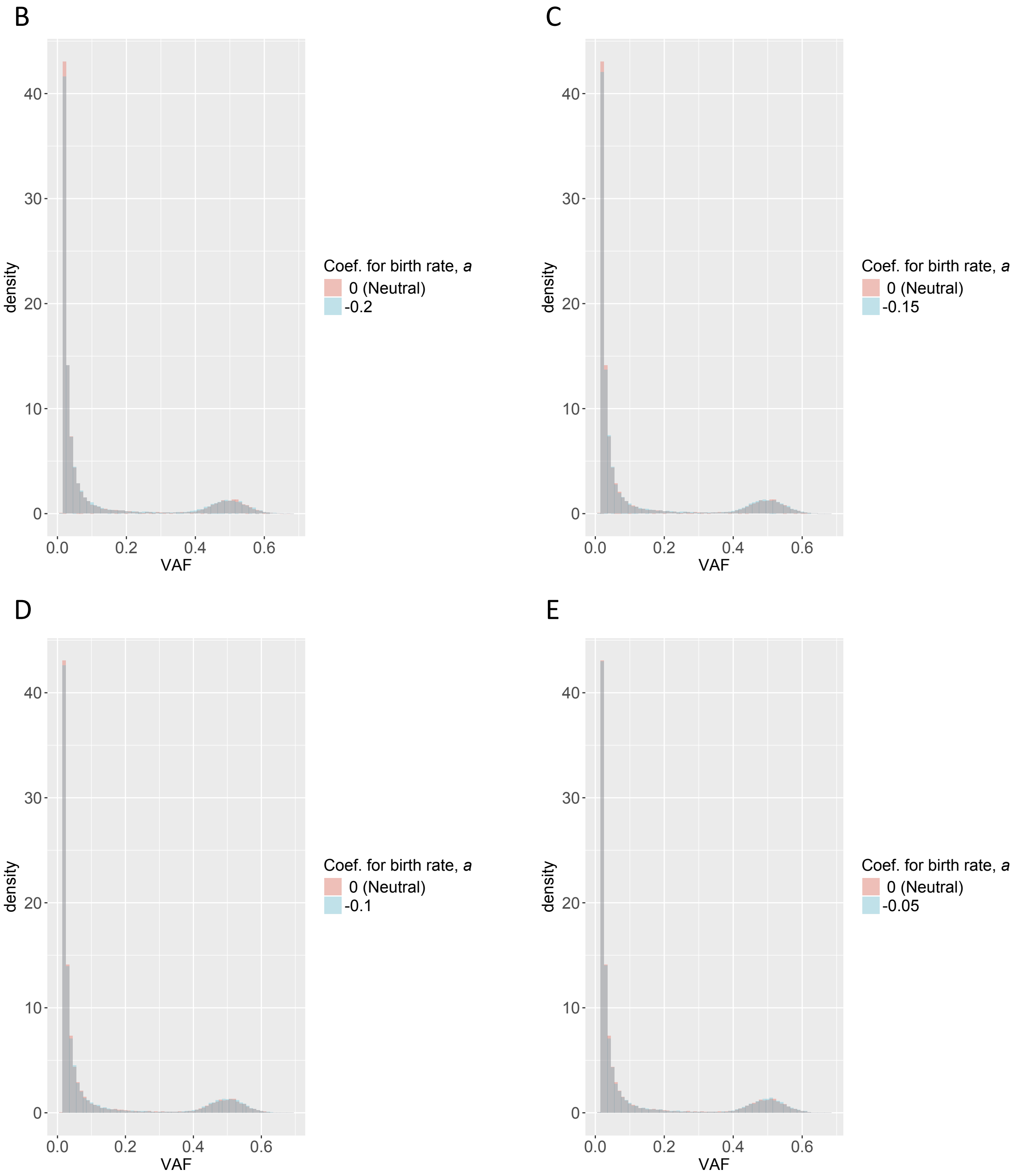

supplementary Figure S13. Continued

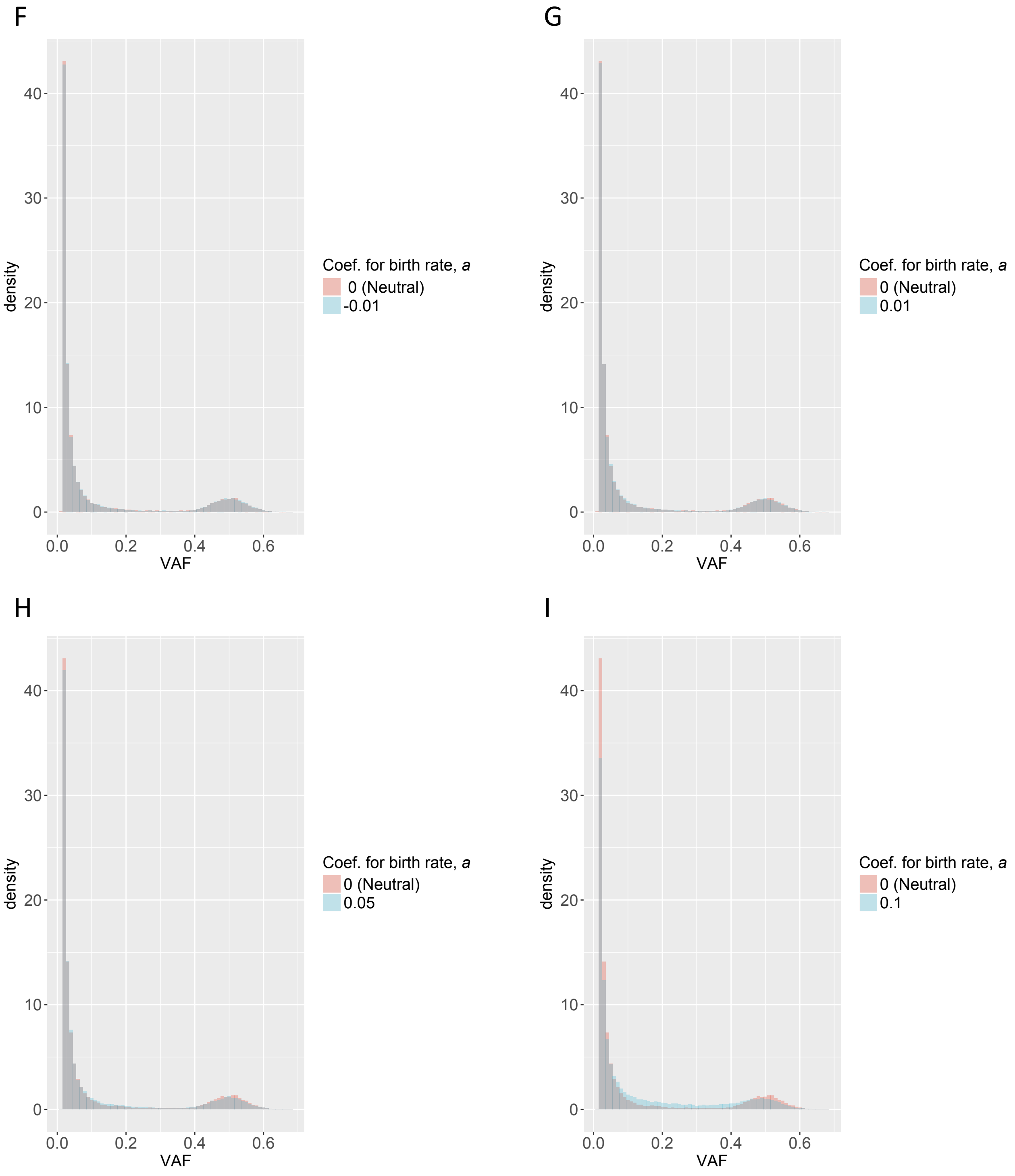

supplementary Figure S13. Continued

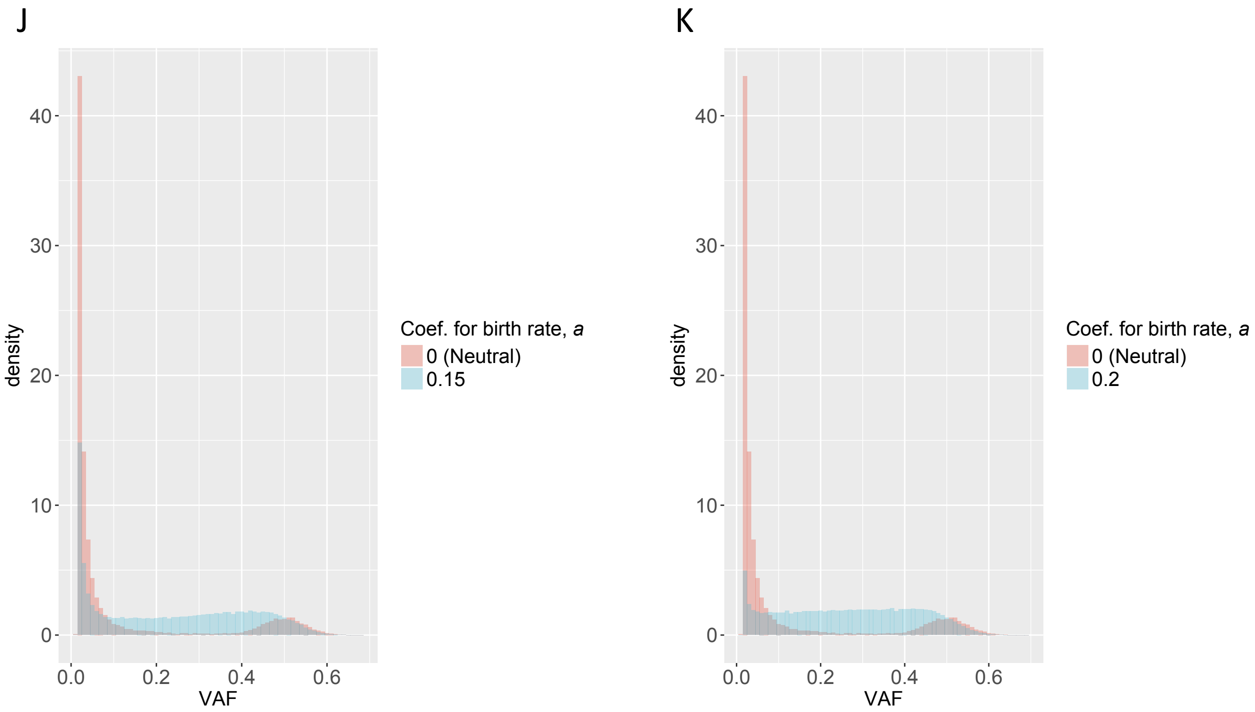

supplementary Figure S14.

A

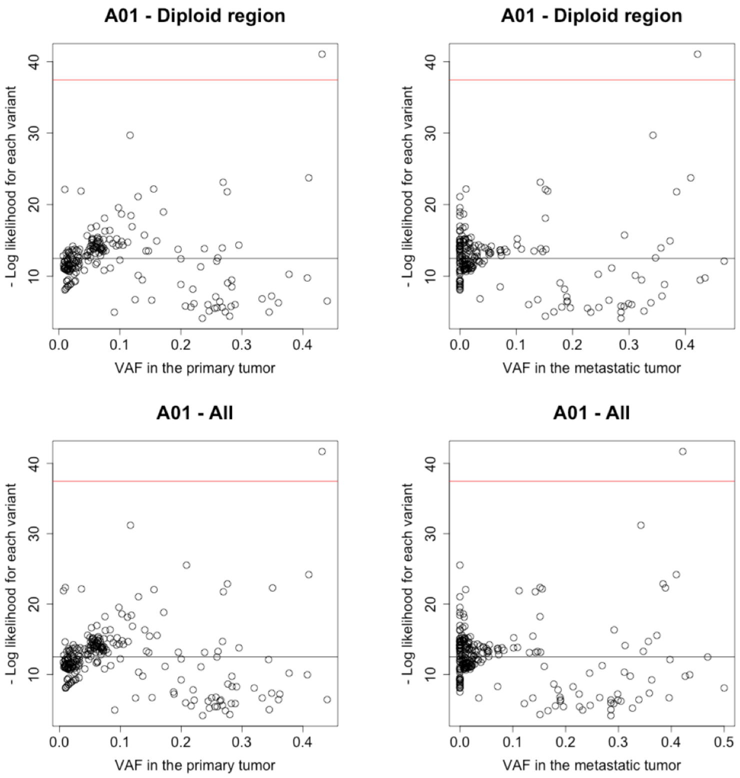


supplementary Figure S14. Outliers detected by preliminary analyses for CRC data of Wei et al. (2017). Black line: Mean of minus log-likelihood for the fitted model for each variant. Red line: Threefold mean of minus log-likelihood and variants over the red lines (outliers) were removed from definitive analyses. (A) Subject A01. (B) Subject A02. (C) Subject A03. (D) Subject A04.

supplementary Figure S14. Continued.

B

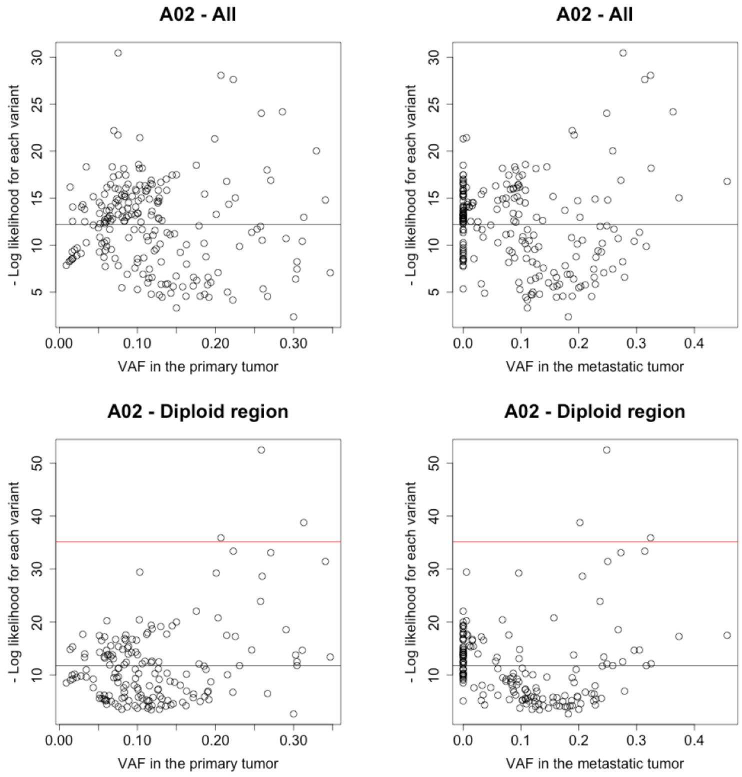

supplementary Figure S14. Continued.

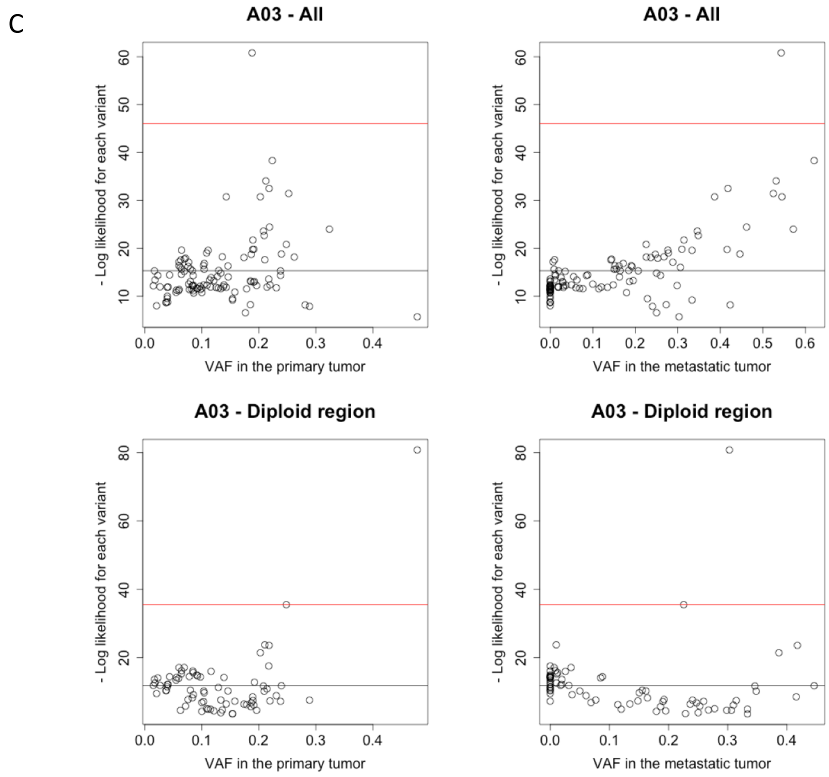

supplementary Figure S14. Continued.

D

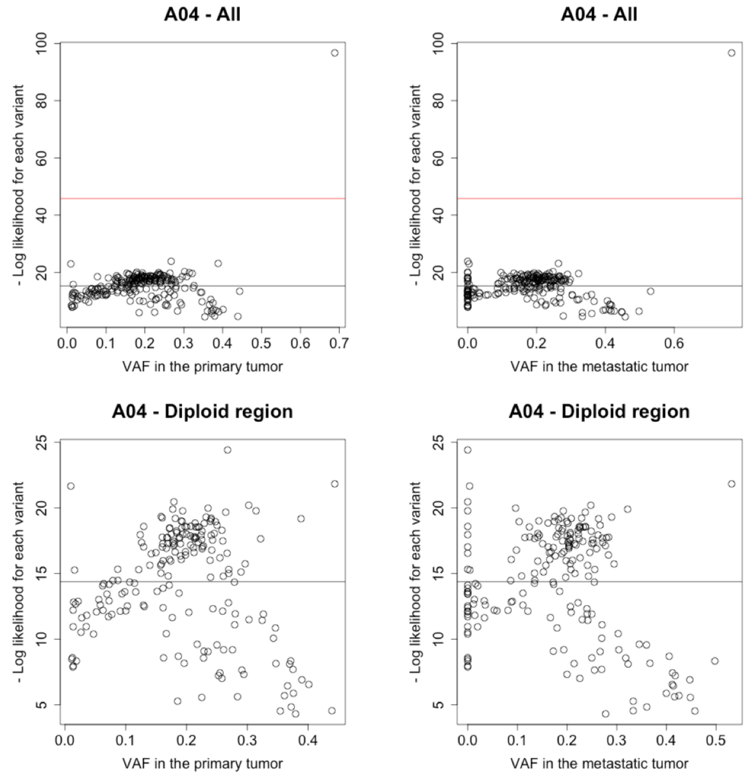

supplementary Figure S15.

A

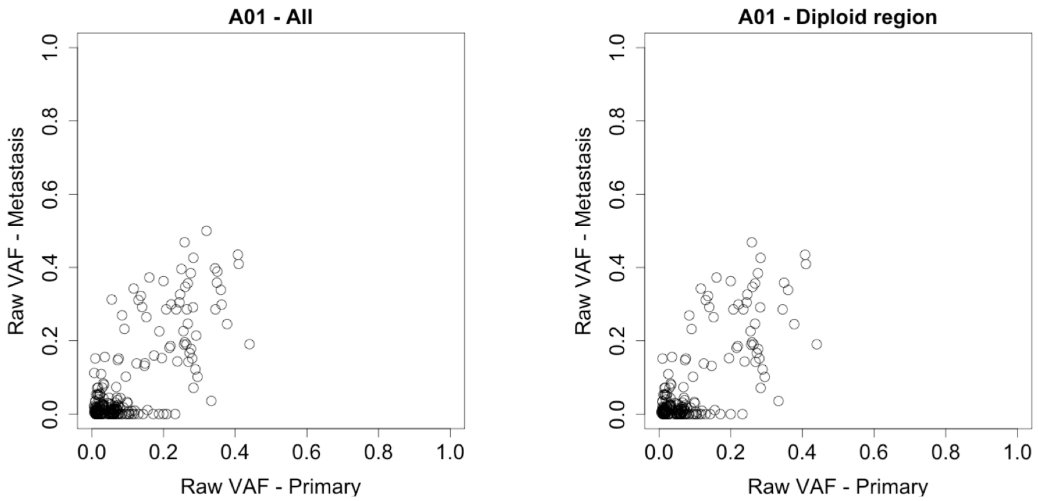


supplementary Figure S15. VAFs of the variants used in the definitive analyses to estimate the founder population size for CRC WES data of Wei et al. (2017). (A) Subject A01. (B) Subject A02. (C) Subject A03. (D) Subject A04.

supplementary Figure S15. Continued

B

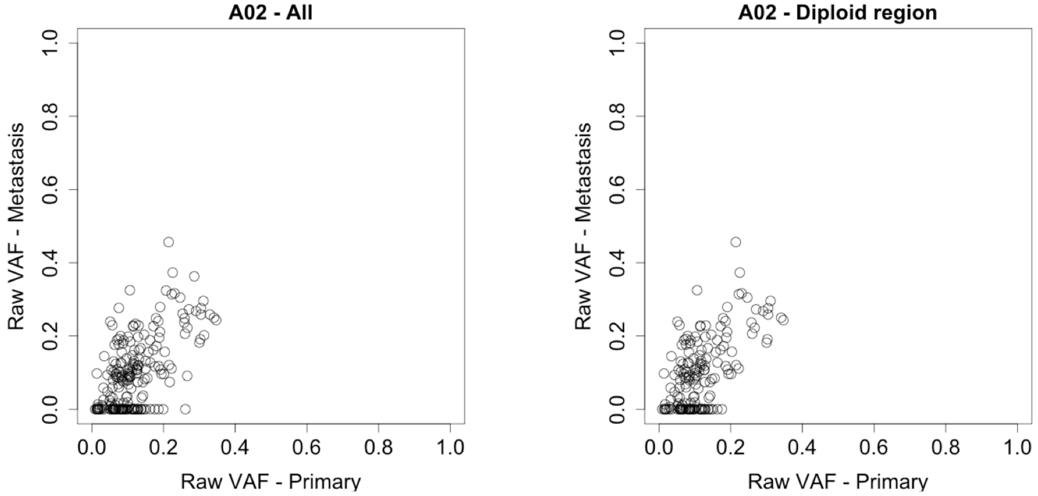

supplementary Figure S15. Continued

C

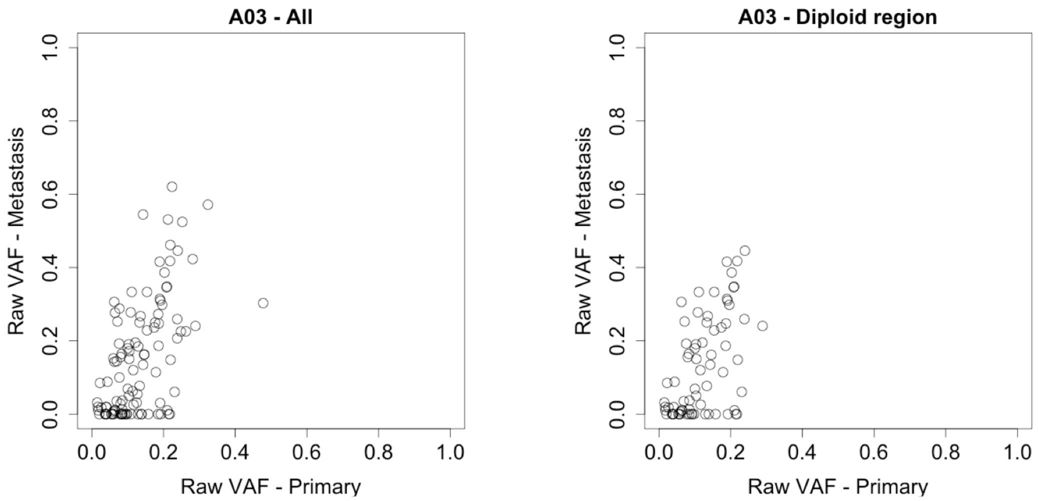

supplementary Figure S15. Continued

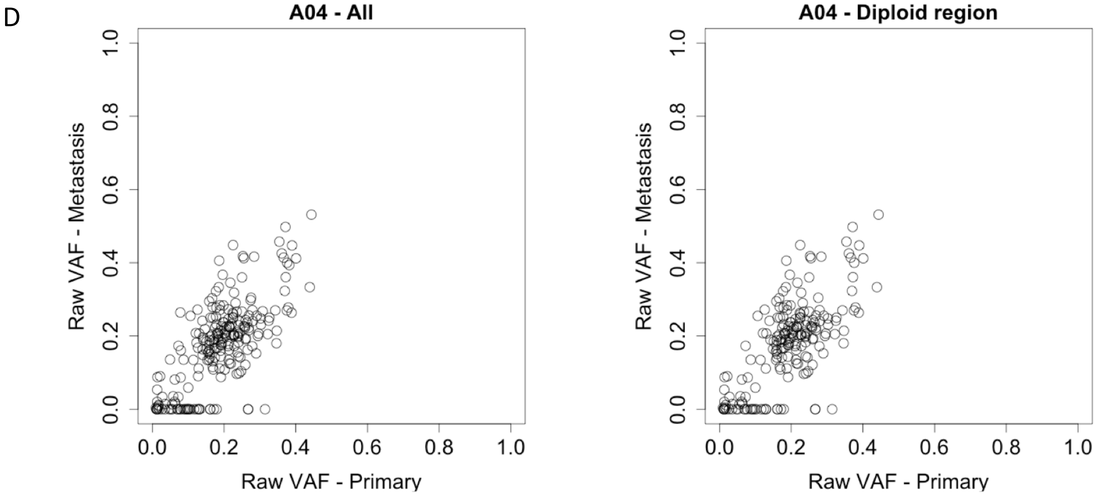
